# Imputing missing counts for waterbird species: A comparison of methods for zero-inflated biodiversity monitoring data

**DOI:** 10.64898/2026.09.24.753523

**Authors:** Barbara Bricout, Laura Dami, Pierre Defos du Rau, Sophie Donnet, Thomas Galewski, Stéphane Robin

**Affiliations:** Sorbonne Université, Université Paris Cité, CNRS, LPSM, 75005 Paris, France; Tour du Valat, Institut de recherche pour la conservation des zones humides méditerranéennes, 13200 Arles, France; Office Français de la Biodiversité, DRAS, Service Conservation et Gestion Durable des Espèces Exploitées, 13200 Arles, France; Université Paris-Saclay, AgroParisTech, INRAE, UMR MIA Paris-Saclay, 91120 Palaiseau, France

**Keywords:** Abundance data, Comparison of imputation methods, Missing data, Population monitoring

## Abstract

1. Species monitoring programmes regularly encounter missing data, which complicates tasks such as estimating population size or detecting temporal trends. Selecting an imputation method suited to the properties of the data is therefore an important practical challenge, particularly for species exhibiting overdispersion and zero-inflation.
2. We compare seventeen imputation methods, comprising thirteen Poisson-based statistical models (accounting for overdispersion and zero-inflation, with fixed, random and multivariate structures) and four contrast-based approaches (LORI, MICE, missForest, correspondence analysis). Using four complete monitoring datasets of waterbird species surveyed across France and Italy over 21 years, illustrating a variety of abundance distributions, we introduced missing observations under a realistic Missing At Random mechanism, at rates from 5% to 70%. We evaluated all methods on computational burden, prediction accuracy for positive and null counts, quantification of uncertainty, and accuracy of population-size estimates.
3. No single method dominates across all criteria. Statistical models and contrast-based approaches yield similar point predictions, but only statistical models provide a genuine measure of uncertainty. Models that jointly account for overdispersion and zero-inflation perform best at predicting zero counts and achieving reliable prediction intervals, and the most suitable method ultimately depends on the abundance distribution of the target species.
4. These results provide practical guidance for ecologists selecting an imputation strategy for incomplete count data, highlighting the trade-offs between predictive accuracy, computational cost, and the ability to propagate uncertainty through subsequent ecological analyses such as population-trend estimation.

## 1 Introduction

Biodiversity monitoring is a cornerstone of ecology and conservation. Large-scale biodiversity surveys are essential for understanding species distributions, population trends, and ecosystem change (Pereira et al., 2013; Yoccoz et al., 2001; Butchart et al., 2010). Many monitoring programs rely on count data, e.g. number of individuals per species, site, or sampling unit collected on a regular time basis. Because of logistical and financial constraints, geopolitical situations, observer variability, or environmental conditions, surveys are frequently subject to missing observations. For example, in the North-African waterbirds population monitoring (UNEP/AEWA, 2018) study presented in Dakki et al. (2021), the proportion of missing observations reaches 60%.

### Incomplete surveys

Missing data raise methodological challenges when willing to analyse the data (Dixneuf et al., 2021; Hossie et al., 2021; Łopucki et al., 2022). Beyond their effects on precision and bias, missing data can fundamentally limit inference by rendering inapplicable a range of statistical methods that require complete observations (Nakagawa and Freckleton, 2008). Furthermore, missing data is a problem when willing to display temporal and spatial patterns of variations (Atkinson et al., 2006; Goluwa Makkala Gunadasa et al., 2026).

A too large proportion of missing data obviously limits our understanding of how biodiversity varies across space and time, and imputation is a way to fill the gaps and construct tools to highlight evolution (Penone et al., 2014). Observation maps provide a direct representation of spatial heterogeneity in ecological metrics such as species richness, species-specific abundance, or occupancy (Hurlbert and Jetz, 2007; Bricout et al., 2026). They are used to identify biodiversity hotspots (Myers et al., 2000), environmental gradients (e.g., latitudinal or elevational), and areas of rarity (McKerrow et al., 2018). As such, they are not merely visual outputs but also core analytical objects in ecological research. These spatial representations benefit from comprehensive spatial coverage at each time to enhance ecological interpretation. Statistical imputation methods offer a rigorous framework to reconstruct complete count data while explicitly accounting for uncertainty, thereby enabling the recovery of more continuous and ecologically meaningful spatial patterns.

Imputation is also useful for the estimation of temporal trends (Haubrock et al., 2025). These trends are essential for tracking biodiversity loss, detecting regime shifts and assessing the effectiveness of conservation actions. Importantly, in this context, the purpose of imputing missing observations is not necessarily to reconstruct individual counts as accurately as possible, but to enable reliable subsequent ecological inference, such as the estimation of annual abundances or population trends. Such analyses similarly depend on complete long-term monitoring series, which are often affected by missing observations. Imputation methods are used to reconstruct complete time series, thereby enabling the estimation of temporal trends while properly propagating uncertainty of prediction.

### Standardized monitoring programs

The present work focuses on standardized biodiversity monitoring programs, rather than opportunistic observation schemes such as citizen-science databases. In standardized surveys, the sampling design, observation protocol, and survey effort are predefined, making the missing-data problem primarily one of incomplete observations rather than heterogeneous sampling effort.

An important feature of these monitoring programs is that they can include the systematic collection of environmental covariates describing the spatial and temporal context of the surveys. These covariates may be site-specific (constant over time), time-specific (common to all sites), or site–time specific. In the latter case, many variables do not require a field visit to be available, as they can be retrieved from external sources such as meteorological or remote-sensing databases (e.g., temperature or precipitation). In this work, we therefore consider the setting in which the abundance observations may be missing, potentially for a large proportion of site–time combinations, while all environmental covariates are assumed to be fully observed. This framework is representative of many long-term standardized monitoring programs and allows us to investigate how environmental information can be exploited to improve the imputation of missing abundance data.

### Missingness mechanism

Although environmental covariates provide valuable information for predicting missing abundances, the validity of the imputation also depends on the mechanism that generated the missing observations underlying the missingness of the observation, as this determines whether the missingness process must be explicitly modelled. Following the classical typology of Rubin (1996) and discussed in Hughes et al. (2019), three missing-data mechanisms are commonly distinguished. Data are Missing Completely At Random (MCAR) when the probability that an abundance is missing is independent of both the observed and unobserved data. In biodiversity monitoring, this would occur typically if the visit of the site is driven by observer availability and unrelated to the ecological phenomenon of interest, to the geographical position or the climate conditions or if a random technical failure results in the loss of a subset of observations. Data are Missing At Random (MAR) when the probability of missingness depends only on observed variables (covariates and observed abundances), for instance if the observation is missing due to the site’s geographical remoteness. Finally, data are Missing Not At Random (MNAR) when the probability of missingness depends on the unobserved abundance itself, even after accounting for the observed variables. For instance, observers may be less likely to report surveys with extremely low abundances, or inaccessible sites may systematically correspond to unusually high or low species abundance. In this case, the missingness mechanism is informative on the missing value and must be modelled jointly with the abundance process.

In standardized monitoring programs, the MAR mechanism arises naturally when survey cancellation is due to variables such as weather conditions, site accessibility, or administrative constraints, that can be reported independently from the visit itself. The covariates of these visits are observed and so can be incorporated into the imputation model. In this paper, we therefore assume that the missingness mechanism is Missing At Random (MAR) and consider imputation methods based on observed abundances and/or available covariates. Under this assumption, the observation process is not considered informative about the underlying ecological processes generating species abundances.

### Imputation methods for biodiversity data

In this paper, we focus on the comparison of imputation methods for incomplete biodiversity count data. Our objective is not to address the broader question of ecological model building, such as the identification and selection of relevant environmental drivers, but rather to assess how different statistical approaches reconstruct missing abundance observations given the available information. Numerous imputation methods have been proposed in the statistical literature (see, for example Grzesiak et al., 2025, for a general overview and an in-depth comparative study). Here, we focus on approaches specifically applicable to ecological abundance data, which must account for specific features of such data, including overdispersion, excess zeros, and dependencies among observations.

Existing methods can be broadly classified into univariate and multivariate approaches (see Table 3 and Sections 2.2-2.3). Univariate methods model each abundance observation separately, typically using (generalized) linear or mixed models. Since these models provide predictions conditional on a set of covariates, they can naturally be applied for imputation when these covariates are available. Examples include TRIM (Van Strien et al., 2004), mixed Poisson regression (Breslow and Clayton, 1993), negative binomial regression (Hilbe, 2011), zero-inflated models (Lambert, 1992), and threshold models (Martin et al., 2005). However, by treating variables independently, these approaches cannot exploit dependencies among observations to improve prediction.

Multivariate approaches aim at exploiting dependencies among observations, either through explicit joint modelling or through data-driven approaches. Among model-based methods, we consider Poisson log-normal models (Aitchison and Ho, 1989; Chiquet et al., 2021) and their zero-inflated extensions (Batardière et al., 2025; Bricout et al., 2026). For approaches that do not rely on an explicit joint probability model, we consider LORI (Robin et al., 2019), MICE (Van Buuren and Groothuis-Oudshoorn, 2011), missForest (Stekhoven, 2025), and correspondence analysis-based methods (Josse and Husson, 2016).

### Comparison of methods

We compare these methods not only in terms of predictive performance, but also according to several methodological criteria. In particular, we examine their underlying statistical assumptions, the data-generating processes they implicitly or explicitly represent, their ability to account for overdispersion and zero inflation, and the criteria used for parameter estimation and optimization. We further investigate how uncertainty associated with imputation can be quantified and propagated, considering, when available, measures such as conditional confidence intervals and asymptotic variance estimates. These aspects are essential in ecological applications, where imputed values are often subsequently used for inference rather than solely for prediction.

The predictive performance of the different approaches is evaluated on a biodiversity datasets issued from the International Waterbird Census (IWC, Wetlands International, 2010), a long-term monitoring program coordinated by [Wetlands International](https://www.wetlands.org/). More precisely, we focus on French and Italian sites surveyed between 1992 and 2012. In this program, the datasets are complete and, for the purposes of this study, we artificially introduce missing observations according to a Missing At Random (MAR) mechanism. This design allows us to compare imputed values with the corresponding true abundances while reproducing a realistic missing-data setting where the probability of missingness depends on available covariates.

All statistical models considered in this work incorporate covariates through a linear predictor. More flexible relationships between environmental variables and abundances could be introduced, for instance, through Generalized Additive Models (GAMs) (Wood, 2017; Smith et al., 2019). However, such extensions primarily concern the specification and selection of ecological covariates rather than the imputation framework itself. Since our objective is to compare imputation strategies given available covariates, we do not consider GAM-based extensions in this study.

### Structure of the paper

In Section 2, we introduce the notations and present the imputation methods considered in this study. We propose an original classification of the methods based on the types of dependencies among observations that they exploit. For each method, we describe the type of inference it enables, including point prediction and the quantification of imputation uncertainty through confidence intervals or variance estimates. In Section 3, we present the Franco-Italian dataset and the in-depth simulation study we designed on it to highlight the respective strengths and limitations of these approaches in various respects.

## 2 Material and Methods

We first introduce the mathematical framework for the data and covariates, together with the notations used throughout this work. We then present the imputation methods under comparison, first considering approaches that assume the counts *Y*_*ij*_ are independent (hereafter referred to as univariate approaches), followed by approaches that account for temporal dependence between counts observed at the same site (hereafter referred to as multivariate approaches).

Table 3 summarizes all the imputation methods considered in the numerical experiments. It highlights their main modelling characteristics and indicates the corresponding R packages available for their implementation.

### 2.1 Data and notations

#### Count matrix

We consider a typical data set collected on *n* sites and along *p* years. Let *Y*_*ij*_ be the number of individuals of the species of interest counted in site *i* (1≤ *i*≤ *n*) on year *j* (1≤ *j* ≤ *p*) and *Y* = {*Y*_*ij*_ : *i* = 1, …, *n, j* = 1, …, *p*}the set of all the counts for the *n* sites and *p* years (which forms an *n* × *p* matrix). In addition, we assume that *Y* is incomplete, meaning for some sites and years the counts are missing. We denote *O* = {(*i, j*) : *Y*_*ij*_ observed} the set of observed observations and by *ℳ* = {(*i, j*) : *Y*_*ij*_ missing} the set of missing observations; we denote by *Y* ^*O*^ and *Y* ^*ℳ*^ the corresponding sets of counts.

#### Covariates

We assume that each site–year pair (*i, j*) is associated with a vector of *d* covariates, which can be decomposed into three groups: *d*_1_ site-specific covariates 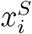, which do not vary over time (e.g., elevation, latitude, or habitat type); *d*_2_ year-specific covariates 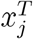, which are common to all sites (e.g., a large-scale climatic index); and *d*_3_ site–year-specific covariates 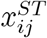, which vary across both space and time (e.g., the temperature or precipitation recorded at site *i* during year *j*). Thus,

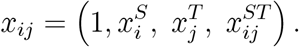

where the one refers to the intercept parameter. We define *X*^*O*^ and *X*^*M*^ as the covariates on the observed and missing counts.

### 2.2 Univariate approaches

A first natural family of approaches consists in exploiting only the information contained in the available covariates, while ignoring possible dependencies among observations. These approaches rely on regression models where observations are assumed to be independent given their covariates, leading to a tractable inference framework. Such regression models are sometimes called species distribution models (SDMs: Elith and Leathwick, 2009). We first introduce the classical Poisson regression model, which provides a natural framework for count data. We then present extensions designed to account for two important features commonly encountered in ecological abundance data: overdispersion and zero inflation.

#### 2.2.1 Poisson regression model

Because we consider counts, the simplest regression model is the Poisson regression model, which states that all counts are independent, with a Poisson distribution, the mean of which depend on the covariates:

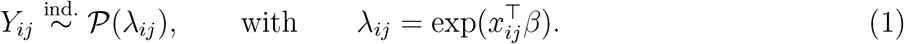

*β* is the vector of regression coefficients encoding the effect of each covariate on the (log-)mean abundance of the species. We consider the Poisson Model (1) as a baseline model. It will be refered as Poisson-FE, for Poisson with Fixed Effects.

#### Remark

Note that the TRIM method (Van Strien et al., 2004) –widely applied for imputation in wildlife monitoring schemes (Lehikoinen et al., 2013; Swaay et al., 2008)– is a particular case of Model (1), including only a site effect and a temporal trend *β*^*T*^ :

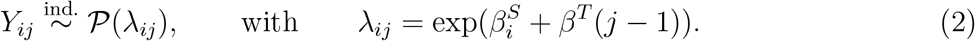

which corresponds to 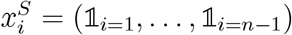. This model will be refereed as Poisson-TRIM in the comparison.

#### 2.2.2 Accounting for over-dispersion

A well-known property of Model (1) is the equality between the mean and the variance: *E* (*Y*_*ij*_) = V (*Y*_*ij*_) = *λ*_*ij*_. This property can be restrictive in many ecological applications, where the variability of V (*Y*_*ij*_) *> E* (*Y*_*ij*_). This departure from the Poisson assumption is referred to as overdispersion (Richards, 2008; Lindén and Mäntyniemi, 2011). A common strategy to account for overdispersion is to introduce additional sources of variability through random effects or latent variables, which increases the variance while preserving the count nature of the observations. Two main choices of latent variable distributions have been considered in the literature to account for overdispersion: a Gamma-distributed latent variable leads to the negative binomial model, while a log-Gaussian latent variable gives rise to the Poisson log-normal model.

#### Negative binomial model

A common choice consists in multiplying the mean *λ*_*ij*_ with a random effect with Gamma distribution with mean one:

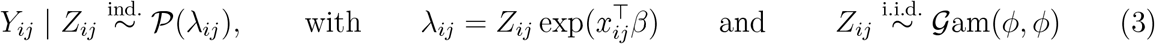

the *Z*_*ij*_’s being i.i.d.; this leads to the negative binomial (NB, Hilbe, 2011) model, refereed to as NB-FE, for Negative Binomial with Fixed Effects in this paper. As shown in Table 1, the additional randomness of the random effect makes the variance of the count larger than its mean, over-dispersion being controlled by *ϕ*.

**Table 1.** Summary of the mean, variance and probability of a zero count for the univariate models. For ZI models, the superscript ‘+’ indicates the conditioning on the presence of the species: 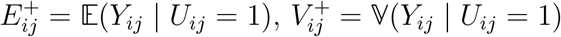 and 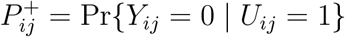, which are given by the corresponding non ZI model.

| Model | $\mathbb{E}(Y_{ij})$ | $\mathbb{V}(Y_{ij})$ | $\Pr\{Y_{ij} = 0\}$ |
| --- | --- | --- | --- |
| (1) Poisson | $\exp(x_{ij}^\top \beta)$ | $\mathbb{E}(Y_{ij})$ | $\exp(-\mathbb{E}(Y_{ij}))$ |
| (3) NB | $\exp(x_{ij}^\top \beta)$ | $\mathbb{E}(Y_{ij}) + \frac{\mathbb{E}(Y_{ij})^2}{\phi}$ | $\left(\frac{\phi}{\phi + \mathbb{E}(Y_{ij})}\right)^\phi$ |
| (4) PLN1 | $\exp\left(x_{ij}^\top \beta + \frac{\sigma^2}{2}\right)$ | $\mathbb{E}(Y_{ij}) + \mathbb{E}(Y_{ij})^2(e^{\sigma^2} - 1)$ | no close form |
| ZI models | $\pi_{ij} E_{ij}^+$ | $\pi_{ij} V_{ij}^+ + \pi_{ij}(1 - \pi_{ij})(E_{ij}^+)^2$ | $(1 - \pi_{ij}) + \pi_{ij} P_{ij}^+$ |

#### Poisson log-normal model

Alternatively, the random effect can be supposed to a have a log-normal distribution, that is:

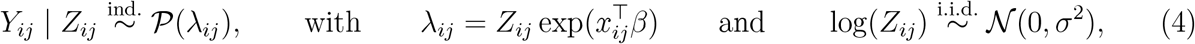

which can be expressed as 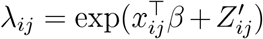 with 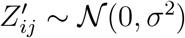. Again the *Z*_*ij*_’s are supposed i.i.d; this corresponds to a generalized linear mixed model (namely, the Poisson mixed model, Breslow and Clayton, 1993), and to the univariate version of the Poisson log-normal model (PLN1: Aitchison and Ho, 1989). Here, over-dispersion is controlled by the variance *σ*^2^ (see Table 1).

#### 2.2.3 Accounting for an excess of zeros

Another specificity of abundance data is the frequent occurrence of zero counts. When the proportion of zeros exceeds what would be expected under standard count models (Equations (1–4)), the data are said to exhibit zero inflation (ZI). An observed zero may have two distinct origins: it may correspond to a true absence of the species at a given site, or to a non-observation despite the species being present, for instance due to imperfect detection. Zero-inflated models (Lambert, 1992; Li et al., 1999) account for these two sources of zeros by introducing a latent binary variable following a Bernoulli distribution.

#### Zero-inflated Poisson model

A first example is the zero-inflated Poisson (ZIP) model, which will be refereed to as ZIP-FE, for ZIP with Fixed Effects, defined as follows:

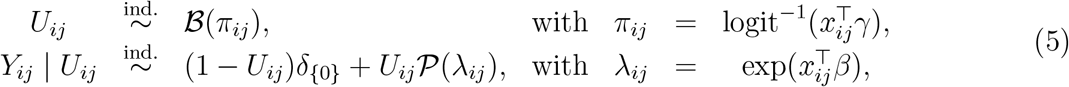

where *δ*{_0_} is the Dirac mass at zero and logit(*p*) = log(*p/*(1 − *p*)). *U*_*ij*_ is the unobserved (or latent) indicator variable for the presence of the species. *γ* is the vector of regression coefficients encoding the effect of each covariate on the presence of the species, whereas *β* encodes their respective effects on its (log-)mean abundance, given its presence. The Dirac mass at zero obviously increases the probability for a null count as compared to the corresponding non-ZI model. Indeed,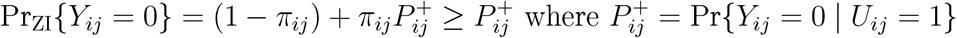.

##### Remark

Note that several formulations (Zeileis et al., 2008) actually model the absence probability, which amounts at changing *U*_*ij*_ into 1- *U*_*ij*_ and switching the sign of *γ*. Also, the logit link function is not mandatory and can be replaced with another function, such as the probit link function: 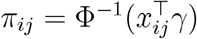, Φ being the cumulative distribution function of the standard normal distribution.

#### Other zero-inflated models

The ZI-NB (later ZINB-FE, for ZI-NB with Fixed Effects) and ZI-PLN1 models are based on the same principle, replacing the *λ*_*ij*_ from (5) with this of (3) and (4), respectively. The mean, variance and probability for a null count for these distributions are given in Table 1.

##### Remark

A distinction must be made between zero-inflated models and hurdle models (Martin et al., 2005): both are based on the same formulation (5) but the hurdle model assumes that zeros can only originate from the latent variable representing presence, so the Poisson part of the model is truncated at zero.

The zero-inflated models ZIP, ZI-NB and ZI-PLN1 can be further enriched with a random term in the Bernoulli part (Hall, 2000; Neelon, 2019):

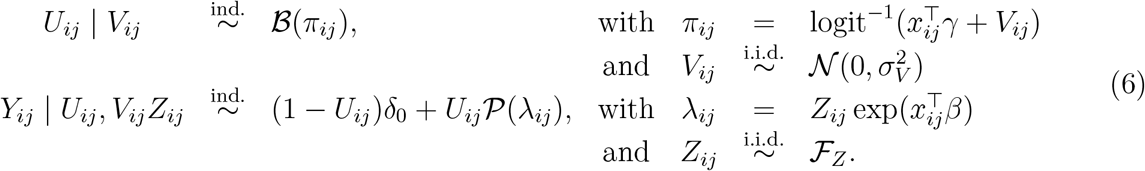

with *ℱ*_*Z*_ = *δ*_1_ for ZIP, or *ℱ*_*Z*_ = *G*am(*ϕ, ϕ*) for ZI-NB or *ℱ*_*Z*_ = Log−*N* for ZI-PLN1. Under Model (6), neither the mean *E* (*Y*_*ij*_) and variance V (*Y*_*ij*_) nor the probability of null count Pr {*Y*_*ij*_ = 0} admit a close form expression.

Univariate approaches are convenient because, thanks to the independence assumption, inference relies only on the observed site-year counts and can be performed using generalized linear mixed models (GLMMs). However, the independence assumption underlying these approaches prevents borrowing information across years when predicting missing abundances. Consequently, the abundance at a given site for a missing year cannot benefit from observations collected at the same site in other years. Improving imputation therefore requires methods that jointly account for observations over time, allowing predictions of missing values to exploit temporal dependence.

### 2.3 Multivariate approaches

We focus here on approaches that share information across years within a given site, but not across sites within a given year (or both). In other words, we do not consider here model involving a spatial dependency structure, but only a time dependency structure. This amounts to assume that the spatial effects are encoded in the covariates or in a random site effect. This choice is motivated by the imputations methods that are actually available at the present time.

We distinguish between two broad families of methods. The first (Subsection 2.3.1) relies on a probabilistic model to capture the dependence structure in the data, whereas the second (Subsection 2.3.2) formulates imputation as an optimization problem by minimizing a suitable loss (or contrast) function.

#### 2.3.1 Probabilistic modeling of dependence

A first approach consist in defining a model for the joint distribution of all the counts observed in a given site, introducing a dependence between all the abundances (observed or missing) from a same site.

##### Generalized mixed linear models

Generalized linear mixed models (GLMM) allow to introduce a random site effect in Model (1), that induces such a dependence structure. Taking

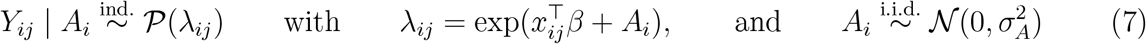

yields the same mean, variance and probability of null count as the PLN1 model in Table 1, but the covariance between two years in a same site is non zero anymore because the random effect *A*_*i*_ affects all the counts *Y*_*ij*_ from site *i*. The covariance between two counts from the same site ℂov(*Y*_*ij*_, *Y*_*ik*_) (*j* = *k*) induced by this random effect is given in Table 2. Under Model (7) (later refereed to as Poisson-RE, for Poisson with Random Effects), counts observed in different sites remain independent so 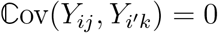 as soon as *i* ≠ *i*^*′*^.

**Table 2.**
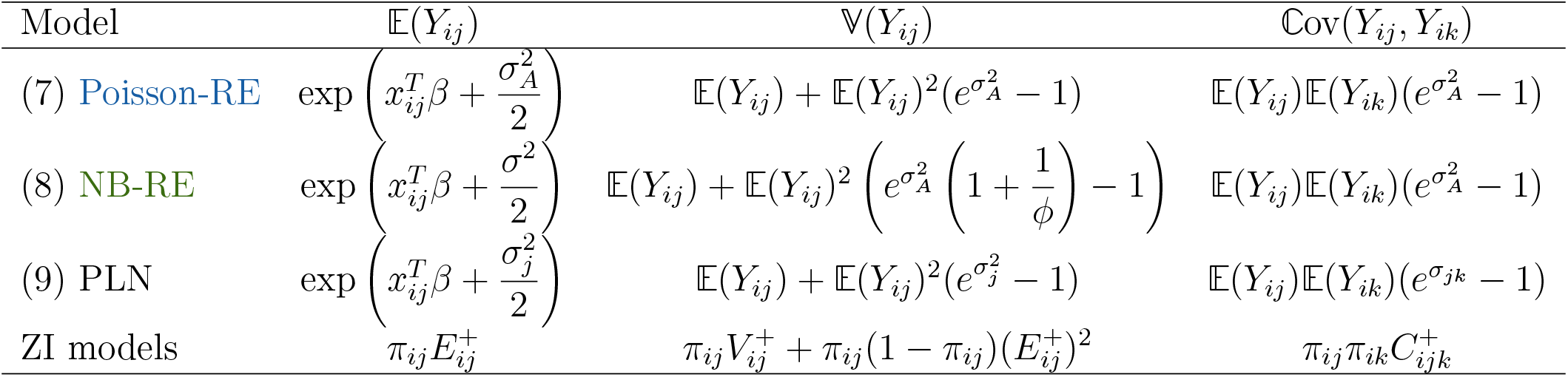
Summary of the mean, variance and covariance for the multivariate models (‘Mixed P’ stands for Mixed Poisson). The covariance ℂ ov(*Y*_*ij*_, *Y*_*ik*_) holds for different years: *j* ≠ *k*. For ZI models, the superscript ‘+’ indicates the conditioning on the presence of the species: 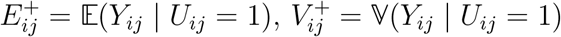 and 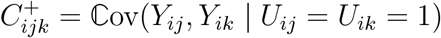, which are given by the corresponding non ZI model.

##### Mixed negative binomial model

A mixed version of the NB model (3), which we will call NB-RE, for NB with Random Effects, including a random site effect can be defined in the same way (Bolker et al., 2009; Stoklosa et al., 2022), taking

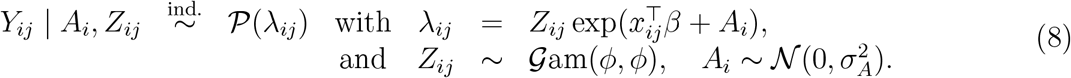

Because of the presence of two random effects (*A*_*i*_ and *Z*_*ij*_) the resulting variance is larger than this of the negative binomial (3), univariate Poisson log-normal (4) and mixed Poisson (7) models (see Table 1), but the covariance is the same as in Model (7) because only the random effect *A*_*i*_ is shared between different years (see Table 2). The probability for a null count has no close form.

##### Multivariate Poisson log-normal model (PLN)

Both the mixed Poisson (7) and mixed NB (8) models yield in a uniform correlation between all years within a given site. A more complex dependency is introduced by the Poisson log-normal model (Aitchison and Ho, 1989), which associates a latent *p*-dimensional random Gaussian vector *A*_*i*_ = [*A*_*i*1_ … *A*_*ip*_]^*⊤*^ with each site:

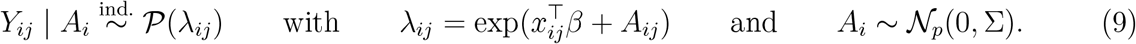

The latent variance matrix Σ = [*σ*_*jk*_]_1_ ≤ _*j*,*k*_ ≤ _*p*_ encodes the dependency between different years via the term *σ*_*jk*_. It also induce over-dispersion via the variance term 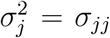 (see Table 2). This model has been proposed by Chiquet et al. (2021) as a versatile joint species distribution model (JSDM).

When dealing with a large number of years, estimating the latent variance matrix Σ proves difficult (Chiquet et al., 2018). A way to circumvent this problem is to assume that the matrix Σ has a low rank *q < p*, that is to assume that there exist a *p* × *q* matrix such that

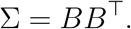

This model can be viewed as a generalization of probabilistic principal component analysis (pPCA: Tipping and Bishop, 1999) to count data and is therefore denoted PLN-PCA. The rank *q* of the covariance matrix Σ can be chosen using a penalized likelihood criterion such as BIC. The presence of a high proportion of missing data further encourages the use of a dimension reduction, and Bricout et al. (2026) recently proposed exploiting a low-rank assumption in this context. From now on, we will call this model ZI-PLN-PCA.

##### Zero-inflated multivariate models

Again, the mixed Poisson, mixed NB (later ZIP-MI and ZINB-MI), and PLN models can be extended to account for an excess of zeros, as in Model (6), by replacing *λ*_*ij*_ with the expressions given in Models (7), (8) and (9), respectively. This extension of the PLN model was proposed by Batardière et al., 2025 and further extended to handle missing data by Bricout et al. (2026) through the ZI-PLN-PCA model. The corresponding mean, variance, and covariance are reported in Table 2, while the probability of observing a zero count is the same as that reported in Table 2 for the corresponding zero-inflated models.

As in the univariate case, these zero-inflated multivariate models can be further extended by introducing a random effect in the Bernoulli component, similarly to Model (6). However, in this case, no closed-form expressions are available for the mean, variance, covariance, or probability of observing a zero count. This leads to the ZIP-RE and ZINB-RE models.

The equations defining all models are provided in Appendix A.1.

#### 2.3.2 Contrast-based approaches

We now present a series of imputation methods that all employ a multivariate approach but do not necessarily rely on an explicit joint distribution.

##### Low-Rank Interaction (LORI)

The `lori` R package (Robin et al., 2019; Dakki et al., 2021) only specifies the form of the expected abundance:

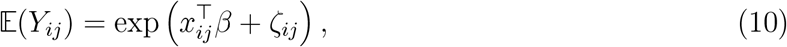

where the parameter *ζ*_*ij*_ encodes potential space-time interactions not included in the covariates vector *x*_*ij*_. Equation (10) is obviously over-parametrized (*np* parameters *ζ*_*ij*_ need be estimated), so the matrix *ζ* = [*ζ*_*ij*_]_1*≤i≤n*,1*≤j≤p*_ is assumed to have a low rank (similarly to the PLN-PCA model).

The parameters *β* and *ζ* are estimated by minimizing a contrast which corresponds to the negative log-likelihood of Model (1) (taking *λ*_*ij*_ = E(*Y*_*ij*_) as given in (10)), to which a penalty term *p*_1_ ∥*ζ* ∥_*\**_ is added to ensure low-rank (∥ · ∥_*\**_ being the nuclear norm). A second penalty *p*_2_ ∥*β* ∥_1_ can also be added to ensure the sparsity of *β*, when dealing with numerous covariates.

##### Multivariate Imputation by Chained Equations (MICE)

A natural way to share information across years within a single site *i* is to model the conditional distribution of the abundance *Y*_*ij*_ given all other years *Y*_*i*,*− j*_ = [*Y*_*i*1_, … *Y*_*i*,*j*_*−*_1_, *Y*_*i*,*j*+1_, … *Y*_*ip*_]^*⊤*^. The R package `mice` (Van Buuren and Groothuis-Oudshoorn, 2011) considers a Gaussian conditional distribution:

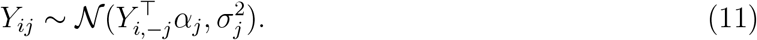

The parameters *α*_*j*_ (1 ≤*j* ≤ *p*) are first estimated by iteratively imputing all observations. Multiple imputation (Rubin, 1987; Rubin, 1996) is then performed by sampling from the set of observed values closest to the predicted value 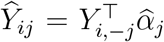. The final step ensures that the imputation is actually a count and provides a sample from the estimated conditional distribution.

##### Random forest (missForest)

The R package `missForest` (Stekhoven, 2025) also relies on the conditional distribution of a given abundance *Y*_*ij*_ given all other abundances from the same site *Y*_*i*,_*−*_*j*_. This approach is nonparametric in that the conditional distribution is estimated using a random forest algorithm. Here again, the algorithm’s parameters are adjusted through iterative imputation of all observations.

##### Correspondence Analysis (CA)

The last approach we considered is based on a classical tool for multivariate analysis, that has been extended to deal with missing data in the `missMDA` R package (Josse and Husson, 2016). CA can be viewed as a dimension reduction technique similar to PCA, but adapted to non-negative data. Denoting *N* = Σ_*i*,*j*_ *Y*_*ij*_ the total count, CA is based on the matrix of relative frequencies *P* = *Y/N*, along with the row and column marginal distributions *r* and *c*. The method consists in performing a singular value decomposition (SVD) of the centered and rescaled matrix

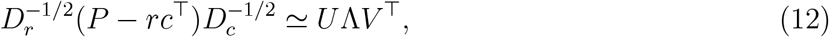

where *D*_*r*_ and *D*_*c*_ are diagonal matrices containing the row and column margins *r* and *c. U* and *V* are orthonormal matrices with dimension *n* ×*q* and *p*× *q*, respectively, and Λ is a diagonal matrix with dimension *q*: hence, CA consists of a low-rank approximation of a modified version of the count matrix *Y*. SVD can not be achieved in presence of missing entries in the matrix *P* but an iterative procedure has been proposed by Josse and Husson (2012) to achieve it in this case. Missing values can then be imputed with the corresponding entries of the reconstructed matrix 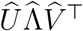. The procedure is implemented in the `imputeCA` function from the `missMDA` package (Josse and Husson, 2016).

Note that, in the last three methods (MICE, missForest and CA), the environmental covariates (*x*_*ij*_) are not used to impute the missing observations. Table 3 summarizes the characteristics of the methods presented in this section. Besides, the last column provides the R package we used in our comparative study. Other R packages (e.g. `glmmADMB, GLMMadaptive, brms, R-INLA`) offer comparable functionalities for fitting Poisson, NB, ZI-Poisson and ZI-NB models with random effects and could be used interchangeably. However, we opted for `glmmTMB` (Brooks et al., 2017) for all such models in this study, for consistency and because of its widespread use in ecology.

**Table 3.** Characteristics of the imputation methods compared in this study. Predictions: ‘marg’ = marginal (13), ‘cond’ = conditional (14), ‘plug’ = plug-in (15) and ‘spec’ = specific to the method.

| | Method | Envir.<br>Covariates | Zero<br>infl. | Multi-<br>variate | Low<br>rank | Over-<br>disp. | CI + PI | Prediction<br>$\hat{Y}_{ij}$ | Package |
| --- | --- | --- | --- | --- | --- | --- | --- | --- | --- |
| Probab.<br>models | Poisson-TRIM (2) |  |  |  |  |  |  | marg | rtrim |
|  | Poisson-FE (1) (17) | ✓ |  |  |  |  | ✓ | marg | stats |
|  | NB-FE (3) (18) | ✓ |  |  |  | ✓ | ✓ | marg | MASS |
|  | Poisson-RE (20) | ✓ |  | ✓ |  | ✓ | ✓ | plug | glmmTMB |
|  | NB-RE (8) (21) | ✓ |  | ✓ |  | ✓ | ✓ | plug | " |
|  | PLN-PCA (19) | ✓ |  | ✓ | ✓ | ✓ | ✓ | cond | colvR |
|  | ZIP-FE (5) (22) | ✓ | ✓ |  |  |  | ✓ | marg | glmmTMB |
|  | ZINB-FE (23) | ✓ | ✓ |  |  | ✓ | ✓ | marg | " |
|  | ZIP-RE (26) | ✓ | ✓ | ✓ |  | ✓ | ✓ | plug | " |
|  | ZINB-RE (27) | ✓ | ✓ | ✓ |  | ✓ | ✓ | plug | " |
|  | ZIP-MI (24) | ✓ | ✓ | ✓ |  | ✓ | ✓ | plug | " |
|  | ZINB-MI (25) | ✓ | ✓ | ✓ |  | ✓ | ✓ | plug | " |
|  | ZI-PLN-PCA (28) | ✓ | ✓ | ✓ | ✓ | ✓ | ✓ | cond | colvR |
| Contrast | LORI (10) | ✓ |  | ✓ | ✓ | Not expl. |  | spec | lori |
|  | MICE (11) |  |  | ✓ |  |  |  | spec | mice |
|  | missForest |  |  | ✓ |  |  |  | spec | missForest |
|  | CA (12) |  |  | ✓ | ✓ |  |  | spec | missMDA |

### 2.4 Imputed value and uncertainty quantification

Given the observations 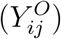 and possibly the covariates 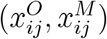, all the methods we presented provide an imputation value 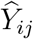 for the missing sites/years. For contrast-based methods such as MICE, MissForest and CA, the imputation calculation do not use the covariates 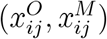 and only rely on 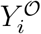 into the evaluation criterion (see, for instance, Equation (11) for the MICE method). We won’t further discuss this task since it as a the core of the methods.

For LORI and model-based methods, the model parameters are estimated by maximizing the (penalized) likelihood or a lower bound of the likelihood when variational inference is used, as in `colvR`- and imputation reduces to a prediction task based on the estimated parameters 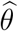. Although this is a classical and seemingly straightforward task which generally relies on an expectation of the form *E* [*Y*_*ij*_| · · ·], there is surprisingly little consensus on which expectation should be used, particularly in multivariate generalized linear models. In the next subsection, we address this fundamental issue by rigorously defining all the prediction forms implemented in the models/packages considered in our comparison. In Subsection 2.4.2, we discuss the possibility to provide a quantification of the uncertainty associated with the imputed value.

#### 2.4.1 Point imputation

LORI and univariate approaches rely exclusively on the covariates associated with the missing observation. The natural imputed value is therefore the *marginal prediction*

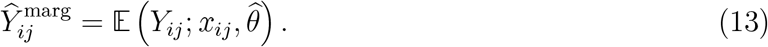

By contrast, multivariate approaches can exploit the dependence with observed abundances from the same site and the natural predictor is then the *conditional prediction*:

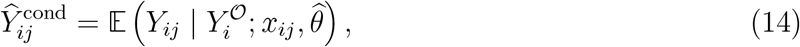

which borrows information from both the available covariates and the observed abundances from the same site. 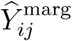 and 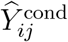 are confounded for univariate methods. In all the multivariate probabilistic models we presented, the dependence relies on a latent variable *ϕ*_*i*_:

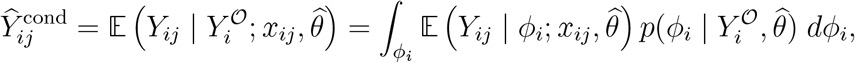

where *ϕ* = *Z*_*i*_ in PLN-PCA (9) and *ϕ* = *A*_*i*_ in the Poisson-RE model (7). Due to its integral form, evaluating the conditional prediction 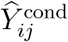 may require non-trivial analytical or numerical computations, and not all packages provide conditional expectations. A simple strategy is to evaluate 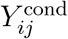 using a Monte Carlo procedure, where the *ϕ*_*i*_ are sampled from 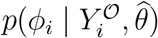. This requires access to, or the ability to sample from, this distribution.

To circumvent this computational difficulty, an alternative prediction is sometimes provided, which we refer to as the *plug-in prediction*, defined as

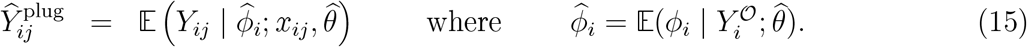

which is more straightforward to compute. Although 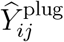 and 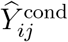 are equal in the case of a linear model –because of the linearity of the expectation– this equality does not hold in general. As an illustration, in the case of the Poisson-RE model (7),which uses the exponential link function, we have

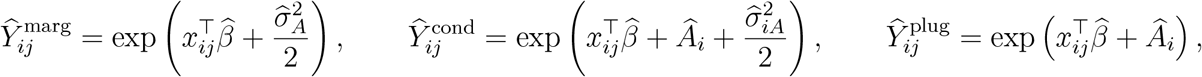

where 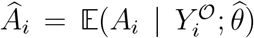 is the estimate conditional expectation of the random effect *A*_*i*_ and 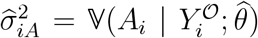 its estimated conditional variance and 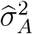 the estimated marginal variance, common to all sites. Notably, in this case, we necessarily have 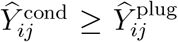, which means that the plug-in estimate 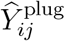 systematically underestimates the conditional mean.

**In practice**, `colvR` provides 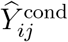 based on a Gaussian approximation of 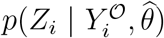 for the ZI-PLN-PCA model. On the contrary, the `glmmTMB` supplies the plugin prediction. The type of prediction provided by each method is summarized in the column ‘Prediction’ of Table 3.

#### 2.4.2 Uncertainty of the imputed value

In addition to an imputed value, it is desirable to provide a measure of the uncertainty associated with it. In practice, few packages provide a direct quantification of the uncertainty associated with individual imputed values. MICE and LORI account for imputation uncertainty through multiple imputation, but do not directly provide prediction intervals for each imputed value. Similarly, `missforest` provides a single imputed dataset together with out-of-bag estimates of imputation error, rather than prediction intervals for individual imputations.

Model-based methods offer a natural framework for quantifying prediction uncertainty. The first source of uncertainty we consider is therefore that associated with the inference procedure. In probabilistic models, this uncertainty is typically quantified through the variance-covariance matrix of the parameter estimator,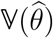. For likelihood-based methods, an estimate of the asymptotic variance, denoted by 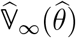, is generally available, yielding the Gaussian approximation

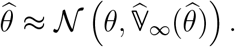

This uncertainty in parameter estimation can then be propagated to the predictions, for instance by constructing confidence intervals for the marginal or conditional expectations defined in Equations (13) and (14).

In general, this propagation can not be done explicitly and we resort to a Monte Carlo procedure where a sample of parameters is generated and the corresponding predictions (marginal, conditional or plugin) are computed

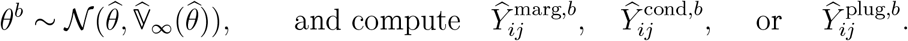

The quantiles of these samples provide *confidence intervals* for the marginal, conditional or plug-in prediction *Ŷ*_*ij*_ which quantifies the uncertainty on the imputed value.

The intrinsic randomness of the observation process can also be be considered. A classical way to account for both sources of uncertainty is to provide a *prediction interval*, that is, an interval (PI) satisfying

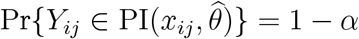

where *α* is a small probability. As before, for a missing observation at site *i* and year *j*, the prediction interval can be defined conditionally on the available information (resulting into the conditional PI):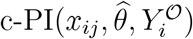. For multivariate approaches, this interval exploits the observed abundances 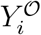 to account for the dependence structure between observations. If ignoring this dependence, one can define the marginal PI m-PI 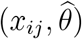. For univariate approaches, c-PI and m-PI are confounded. For multivariate approaches, we expect the conditional intervals to be narrower than the marginal intervals. Again, this double source of randomness is difficult to propagate in a closed form and it is standard to resort to a Monte Carlo procedure:

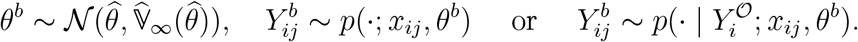

where 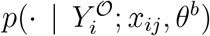 and *p*(·; *x*_*ij*_, *θ*^*b*^) involve latent variables. The quantiles of these samples provide the marginal and conditional prediction intervals. Examples of Monte Carlo procedures to obtain confidence and prediction intervals and provided in Appendix A.3, Algorithms 1 and 2.

##### In practice

only a few packages provide uncertainty measures for predictions. To the best of our knowledge, `colvR` is the only package that directly implements the procedures described above for computing prediction intervals for the ZI-PLN-PCA model. For other models, the same Monte Carlo strategy can in principle be applied provided that the package supplies an estimate of the asymptotic covariance matrix 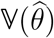. This is the case for `glmmTMB`, which provides such an estimate, but not for `rtrim`. In fact, `rtrim` focuses on estimating population size and trends and provides confidence intervals in which the variance has been propagated, but it does not provide access to this variance nor does it provide prediction intervals for the imputed data.

*Remark*. Temporal trend estimation

Estimating temporal trends is not the main focus of this article, but it is central to conservation work (Wauchope et al., 2019; Rodriguez-Caro et al., 2024; Rueda-Cediel et al., 2015). In particular, it is often made more difficult by missing data (Godeau et al., 2026; Bowler et al., 2025; Łopucki et al., 2022; Dakki et al., 2021). Thus, when there is missing data, there are two ways of approaching the issue of trend estimation. One can impute the missing data and then estimate a trend from the completed data - this is what contrast-based approaches propose - or one can rely on the estimated parameters of a model, together with their variance, to estimate the trend - this is what statistical models propose. The advantage of the second option is that the variance of the estimated parameters can be propagated to the trend, providing a genuine measure of uncertainty around it, whereas a trend estimated from imputed data hides the uncertainty of the imputation step itself. Failing to take this uncertainty into account means running the risk of obtaining a non-significant trend over time, that is to say, one whose direction is uncertain.

## 3 Results

The aim of this section is to compare the imputation methods across multiple dimensions. More specifically, we want to assess which methods provide the most accurate predictions of missing abundances, together with reliable measures of uncertainty for these predictions. To this aim, we considered datasets from four *complete* monitoring programs, from which we removed an increasing proportion of observations. We then assessed the performances of the different methods according to a series of observable criteria, that is criteria that can be evaluated based on the actual abundances and not, for example, their expectations. We also paid attention to the computational time required by each method, and to the proportion of cases where implementation problems (numerical issues, convergence failure, etc.) were encountered by each of them.

### 3.1 Simulation design

#### 3.1.1 Testing datasets

The four complete datasets we use correspond to winter counts of four species of water birds: the Greylag Goose (*Anser anser*), the Northern Shoveler (*Spatula clypeata*), the Mute Swan (*Cygnus olor*), and the Eurasian Coot (*Fulica atra*). The monitoring program took place in France and Italy over *p* = 21 years period, from 1992 to 2012.

These species display very different distributions in this region (see Figure 1). These contrasting distributions provide a useful range of ecological situations for evaluating the methods, from species with relatively sparse occurrence and a high proportion of zero counts to species occurring more consistently across monitored sites and sometimes in large aggregations. Within the datasets considered here, the Greylag Goose and Northern Shoveler show lower occurrence and abundance, whereas Mute Swan and Eurasian Coot are more frequently recorded and generally occur at higher abundances.

**Figure 1.**
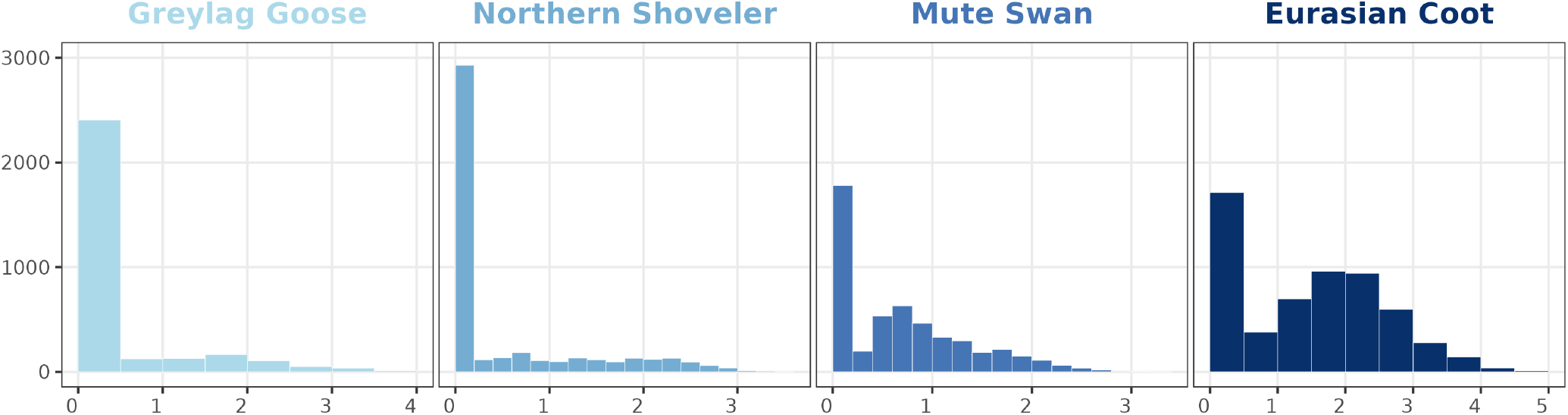
Distribution of log_10_(1 + abundance) of four species of waterbirds in France and Italy : the Greylag Goose (top left, 144 sites), the Northern Shoveler (top right, 214 sites), the Mute Swan (bottom left, 239 sites) and the Eurasian Coot (bottom right, 274 sites).

In addition to the count of each of the four species, we have a covariate describing winter rainfall for each site in each year, with no missing data. In Model (1), we can see that it is also possible to include covariates relating solely to sites or to years; in our case, to achieve greater accuracy (Bricout et al., 2026), we chose to replace such covariates with two generic fixed effects: a site-specific effect and a year-specific effect. Denoting by 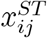 this site-year-specific covariate (winter precipitation)for site *i* on year *j*, the linear term from Model (1) writes

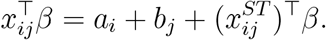

#### 3.1.2 Missingness pattern

We consider a Missing at Random (MAR) scenario of missing data, meaning that the probability of missingness depends jointly on sites and years. This scenario reflects monitoring programmes in which survey coverage varies both among sites and years, for example because some sites are more difficult to survey regularly, while overall monitoring effort has grown over time. We removed a proportion of 5%, 30%, 50%, and 70% of the observations from the original dataset. 100 replicates were generated for each configuration, resulting in 4 × 100 = 400 data sets per species. We denote by *ℳ* the set of index pairs (*i, j*) such that *y*_*ij*_ is missing, and let *M* = card(*ℳ*).

#### 3.1.3 Inference & prediction (inc. zeros)

For each dataset, we applied each of the methods listed in Table 3 using the indicated R package. Once fitted, we computed the prediction for each missing value using the R function `predict` for all the methods implemented with `glmmTMB` and the prediction outputs for the other methods. Depending on the method, *Ŷ*_*ij*_ refers to 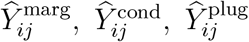, or any other value specific to the method, as specified in Table 3. To better evaluate the benefits of zero-inflated models, we also considered the prediction of null counts. More specifically, for each model, we computed the probability Pr {*Y*_*ij*_ = 0}, which is the probability for the species either to be absent or to be present but not observed, and that can be compared with actual observed count.

In addition to the predictions, for each method providing an estimate of the variance of the parameter estimators 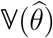, we computed prediction intervals.

#### 3.1.4 Evaluation criteria

We compared the different methods in terms of computational burden, accuracy of their prediction, uncertainty of these predictions. We also focused on their ability to predict null abundances.

##### Computational burden and success rate

We recorded the average computation times as well as the proportion of simulations for which the estimation algorithm associated with each method did actually converge and/or produce an estimation of the variance of the estimators.

##### Prediction accuracy

We first evaluate the precision of the prediction in terms of root mean squared error (RMSE) :

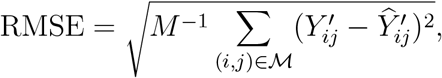

considering three transformations 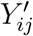 of the count *Y*_*ij*_: the count itself 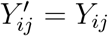, its logarithm (plus one): 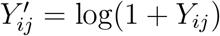 and its square-root: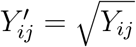.

We will also compute the relative RMSE (RRMSE):

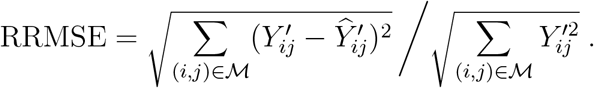

RMSE and RRMSE (without logarithm or square-root transformation) are criteria that specifically penalise large errors, whilst the logarithm and square-root transformations take account of the fact that the variance – and therefore the uncertainty of the errors – depends on the true value.

What interests us is the ranking of the methods according to the criterion, not the value of the criterion itself. However, in this case, we note that RRMSE and RMSE are equivalent, so we have chosen to present only the RRMSE, which is used more frequently (Dakki et al., 2021; Godeau et al., 2026).

##### Measure of uncertainty

For each method providing an estimate 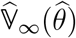 of the parameter estimates, we computed the marginal, conditional or plug-in prediction intervals (depending on the model) at the 90% level. We assessed whether these intervals achieved the nominal coverage rate by comparing their empirical coverage to the 2.5% and 97.5% quantiles of *ℬ* (*M*, 0.9). Finally, we computed the width of the prediction intervals.

##### Prediction of zero

Regarding the prediction of zero counts, missForest and MICE can directly impute a zero count. Neither CA nor LORI explicitly predicts zero counts. In contrast, probabilistic models based on the Poisson or zero-inflated Poisson (ZIP) distributions can provide an estimate of the probability that a missing observation corresponds to a zero count, that is 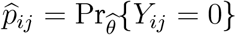. For the latter, we used a threshold at 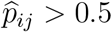 to predict a zero.

This made, we then treated the problem as classification problem comparing the actual *Y*_*ij*_ = 0 or not with the prediction (zero or not) with two standard criteria:

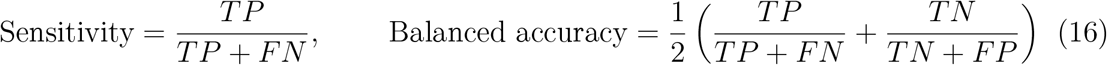

where *TP* stands for the number of correctly predicted zeros, *FP* for the number of falsely predicted zeros, *FN* for the number of actual zeros predicted as non-zeros and *TN* for the number of actual non-zeros correctly predicted as non-zero. For the statistical models, we also computed the ROC curve based on 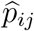 and the corresponding area under the curve (AUC).

### 3.2 Results

#### 3.2.1 Processing time and convergence issues

##### Computational time

Figure 2 displays the computation times for each method on every configuration (species and missing rate). Most of the methods take less than a minute to run. However, it is noticeable that GLMMs with zero-inflation take longer to compute than the others, which is consistent with the fact that zero-inflation involves twice as many parameters to estimate. In contrast, three methods stand out for their substantially longer computation times: PLN-PCA, ZI-PLN-PCA, and LORI. For these methods, a parameter selection step - namely the selection of the latent dimension *q* for PLN-PCA and ZI-PLN-PCA, and a cross-validation procedure for LORI — adds considerable overhead. LORI is the most computationally demanding, with average computation times reaching up to more than twenty minutes. It is worth noting, however, that the data was collected over a period of 21 years and that, on that scale, even 20 minutes is worth waiting for.

**Figure 2.**
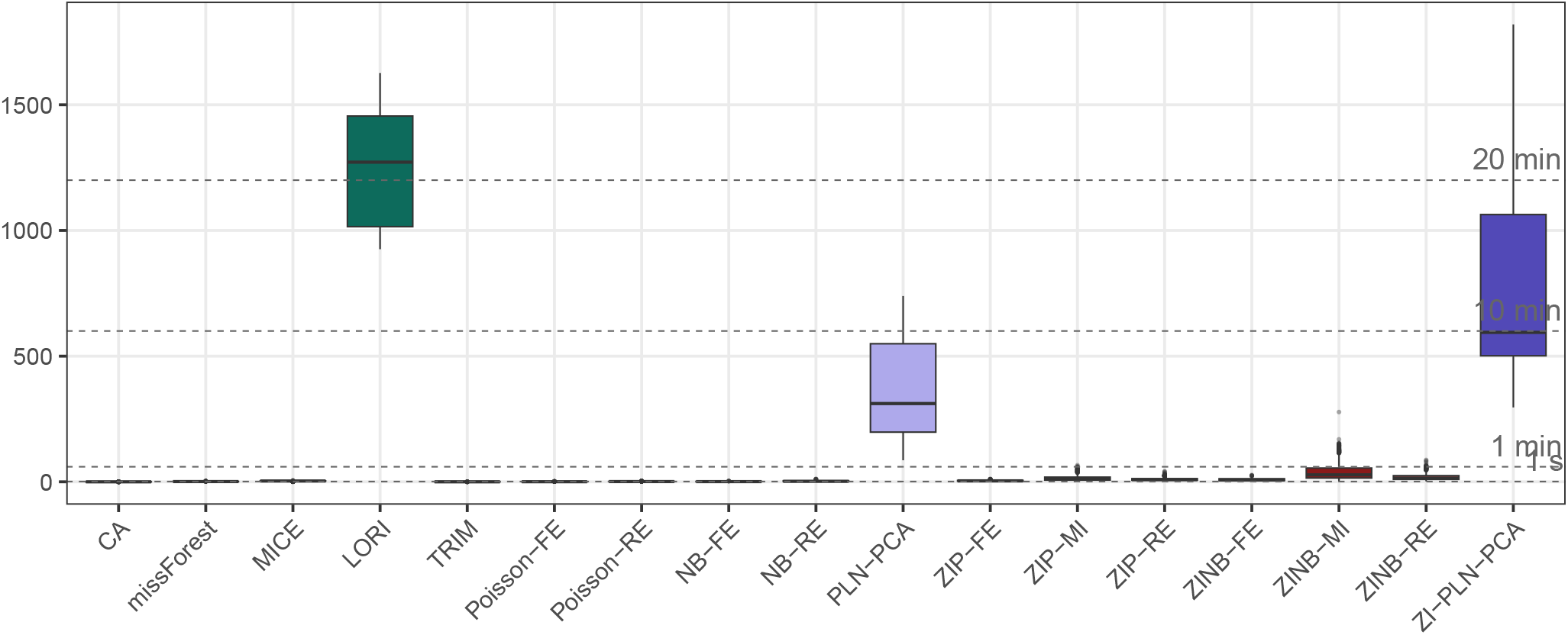
Computation times for each imputation method across all species and missing rates. Times include both the parameter selection step (latent dimension *q* for ZI-PLN-PCA and PLN-PCA, cross-validation for LORI) and the model fitting step.

##### Convergence issues

Across all species and missing rates, all methods successfully produced complete imputations for every replicate, with one exception: MICE. MICE completes 100% of replicates when 30% or less of the data is missing, but its performance degrades sharply beyond this threshold: at 50% missingness, a non-negligible fraction of replicates yield incomplete imputations, and at 70%, MICE fails to produce any complete replicate.

As for the GLMMs, the estimation of the asymptotic variance matrix 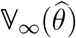 also turned out to fail (the vcov function returning NAs/NaNs) for some simulations. The success rates for the variance-covariance matrix of the estimators are reported in Table 4. Whilst Poisson and Negative Binomial models achieve near-perfect success rates, their zero-inflated counterparts - ZIP-FE, ZIP-MI, ZIP-RE, ZINB-FE, and ZINB-MI-exhibit substantially lower rates. ZINB-RE is a notable exception among zero-inflated GLMMs, suggesting that the combination of a random effect with a negative binomial distribution provides a more stable optimisation landscape. Note that ZI-PLN-PCA relies on a different package and a distinct inference algorithm which enables it to always return a matrix 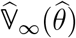.

**Table 4.**
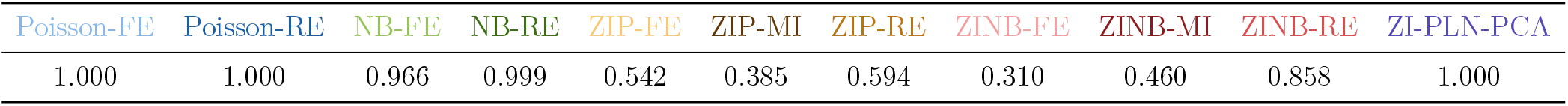
Success rate of the variance-covariance matrix computation with the R function `vcov` for the parameter estimators, pooled across all species and all missing data rates.

| Poisson-FE | Poisson-RE | NB-FE | NB-RE | ZIP-FE | ZIP-MI | ZIP-RE | ZINB-FE | ZINB-MI | ZINB-RE | ZI-PLN-PCA |
| --- | --- | --- | --- | --- | --- | --- | --- | --- | --- | --- |
| 1.000 | 1.000 | 0.966 | 0.999 | 0.542 | 0.385 | 0.594 | 0.310 | 0.460 | 0.858 | 1.000 |

#### 3.2.2 Quality of imputation

##### Imputation errors

In this section, we present the results for the Northern Shoveler with 30% of the entries missing. Similar results were obtained with the other species (not shown) and the other missing rates (shown in Appendix A.2).

The central focus of this article is a comparison of the quality of the imputations produced by the various methods. Intuitively, this simply amounts at comparing the prediction errors of each method, but the definition of the measure of these errors itself is a matter of debate. Indeed, the popular RRMSE criterion is designed to heavily penalise large errors. However, in the case of Poisson-based models, the variance – and therefore the uncertainty of the prediction – depends on the expected value and thus on the missing value itself. A very large count will have greater uncertainty than a small count. Consequently, the error in a large count contributes disproportionately to the calculation of the RRMSE. We can therefore propose RMSE values with a logarithmic or square-root transformation as a criterion that takes into account the proportionality between the true value and the error. The RRMSE favours methods where the error does not necessarily depend on the missing value, which means that the error can be of the same magnitude for both small and large counts, whereas logarithmic and square-root transformations favour methods where the error depends on the true value.

Furthermore, the square root transformation is known to stabilize the variance of the Poisson model (17) (Bartlett, 1936; Anscombe, 1948) and the logarithmic transformation plays an analogous role for the PLN-based models. This is reflected in the results of Figure 3: the square-root RMSE tends to favour Poisson-based GLMMs, while the log RMSE tends to favour PLN-based models. We therefore conclude that three different criteria yield three different rankings of the methods. However, for all criteria, we find that the zero-inflated version of all models (Poisson, Negative Binomial, PLN-PCA) produces smaller errors than the non-zero-inflated version.

**Figure 3.**
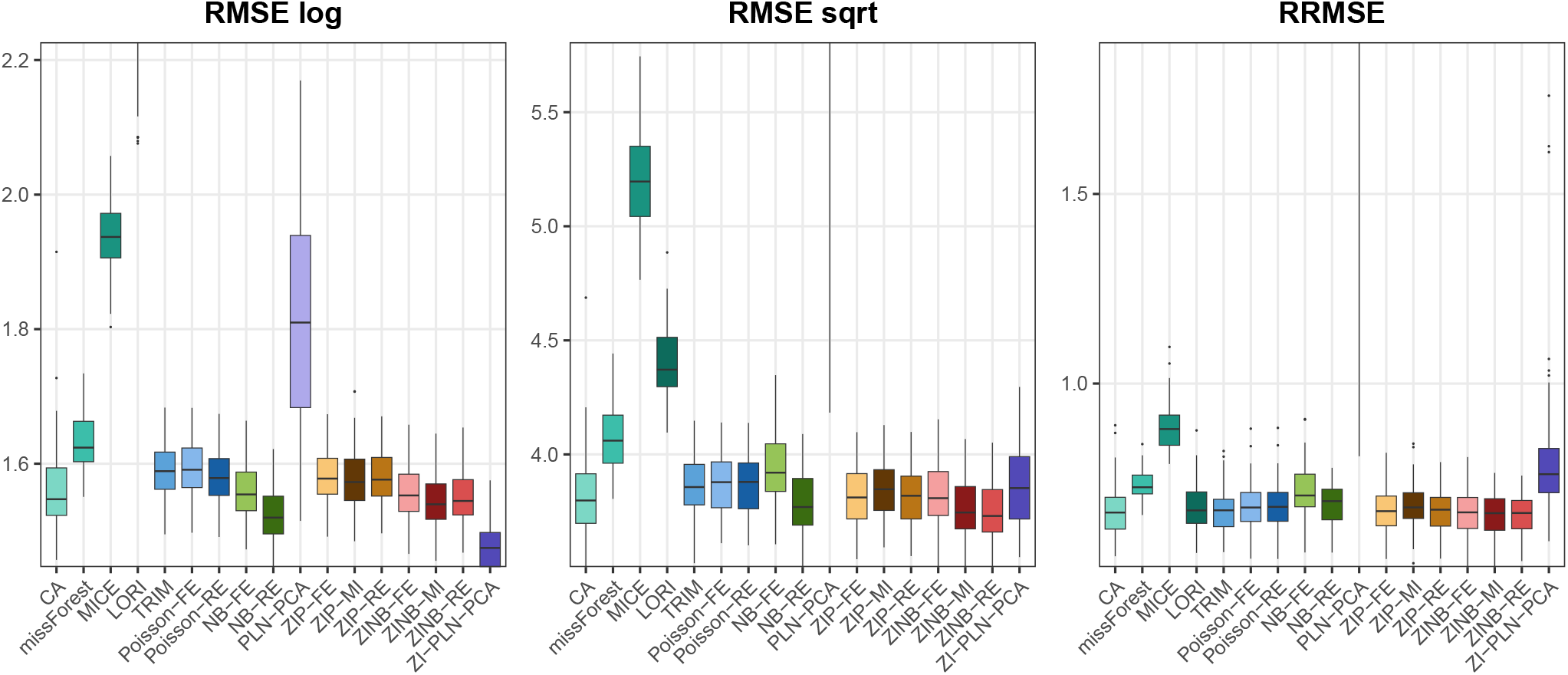
Imputation errors with 30% missing data for the Northern Shoveler for all methods measured using three different metrics.”

To gain a clearer picture of the estimates, we compared them directly with the true values and present these results in Figure 4. The scatter plots reveal that most methods agree on the predictions and align with the truth. This is the case for all GLMs and GLMMs, as well as ZIPLN-PCA and Poisson-TRIM, with the sole exception of PLN-PCA. This suggests that, in terms of mean, these methods are broadly equivalent to each other for this dataset and configuration.

**Figure 4.**
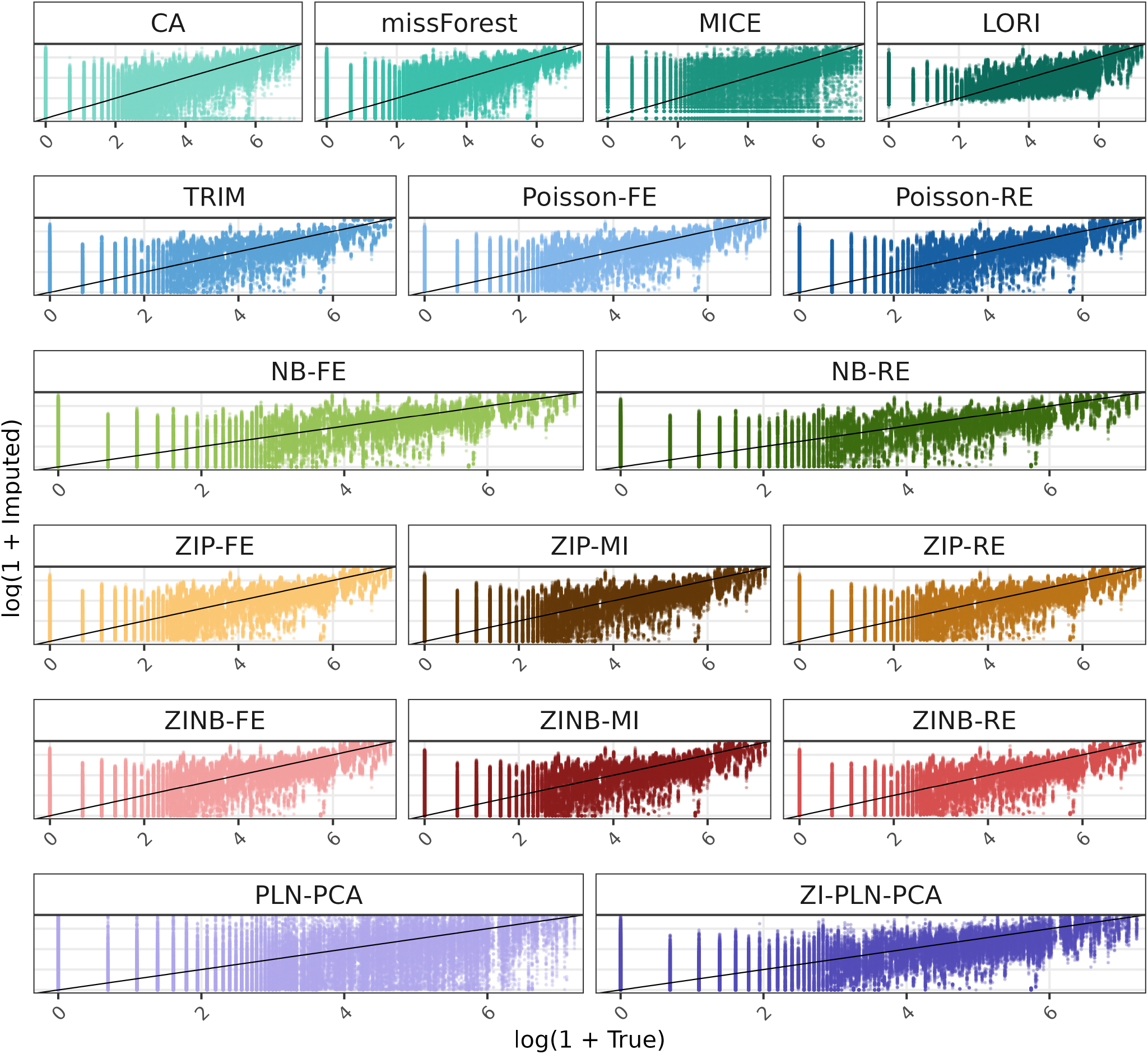
log(1 + Imputed values) for the Northern Shoveler with 30% missing data across all replicates versus log(1 + truth) for each method.

For the other methods, missForest produces predictions similar to those of GLMMs. LORI exhibits a different error pattern: unlike the other methods, it does not predict small values. This explains why the log transformation, which penalises errors on small values more heavily, does not favour LORI. MICE is the method whose imputations differ most from those of the other methods. Therefore, by relying solely on point-by-point predictions, it is not possible to identify one method as being more accurate than the others. In particular, as the nature of the predictions differs depending on the method, it is not even possible to choose a single criterion for ranking the various methods.

##### Prediction intervals

As exposed in Table 3, for all model-based approaches, each prediction can be accompanied with a prediction interval. There are two questions that help to assess the quality of prediction intervals: does the interval adequately cover the true (but removed) value, at the nominal rate, and is the interval narrow enough to be informative?

For each species, for each missing data rate, we took 10 replicates and calculated prediction intervals for the missing data in these 10 replicates. Table 5 shows the average coverage rate per species, taking all missing data rates into account, and indicates with a ✓whether, for the majority of replicates, the results fall within the corresponding binomial bands (*ℬ* (*M*, 0.9) where *M* is the number of missing cells). We also show the width of the prediction intervals by species (for all missing data rates combined) in Figure 5. Note that in these two figures, not all models and details of missing data rates are shown, but they are available in Appendix A.3. We would like to point out that we are comparing prediction intervals which are fundamentally different in nature, as the ZI-PLN-PCA intervals are conditional and those of the GLMM are plug-in.

**Table 5.**
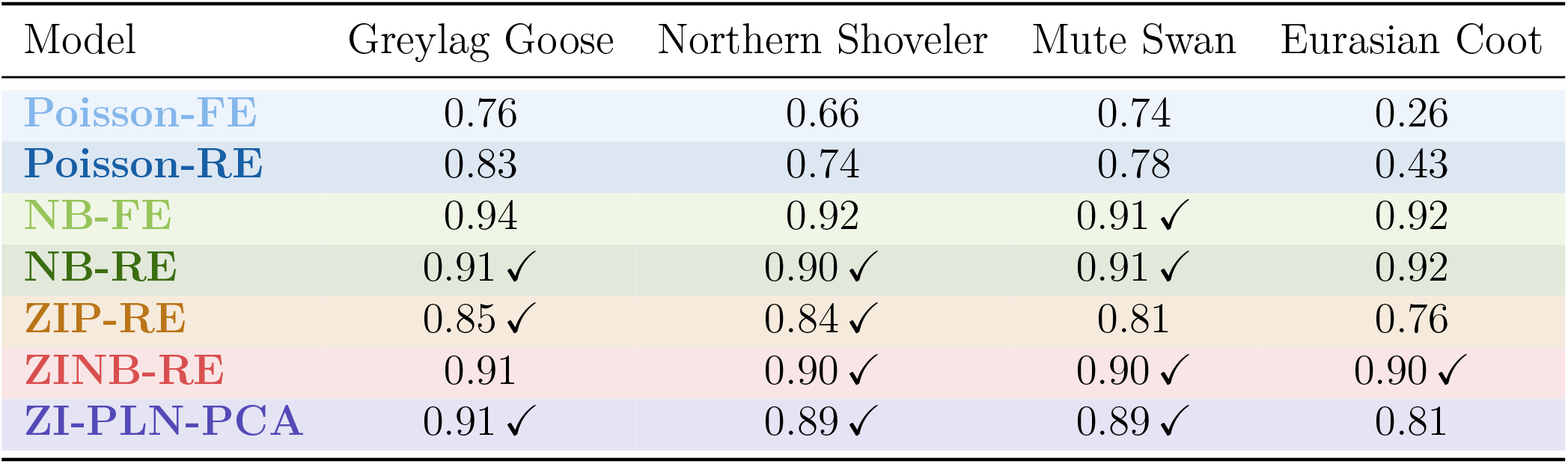
Mean empirical coverage rate, pooled across the four missing-data rates (5%, 30%, 50%, 70%) for the four species. A ✓indicates that at least half of the pooled individual replicates have a coverage rate falling within their own binomial reference band Binomial(*M*, 0.9) (essentially [0.88, 0.91] throughout).

| Model | Greylag Goose | Northern Shoveler | Mute Swan | Eurasian Coot |
| --- | --- | --- | --- | --- |
| **Poisson-FE** | 0.76 | 0.66 | 0.74 | 0.26 |
| **Poisson-RE** | 0.83 | 0.74 | 0.78 | 0.43 |
| **NB-FE** | 0.94 | 0.92 | 0.91 ✓ | 0.92 |
| **NB-RE** | 0.91 ✓ | 0.90 ✓ | 0.91 ✓ | 0.92 |
| **ZIP-RE** | 0.85 ✓ | 0.84 ✓ | 0.81 | 0.76 |
| **ZINB-RE** | 0.91 | 0.90 ✓ | 0.90 ✓ | 0.90 ✓ |
| **ZI-PLN-PCA** | 0.91 ✓ | 0.89 ✓ | 0.89 ✓ | 0.81 |

**Figure 5.**
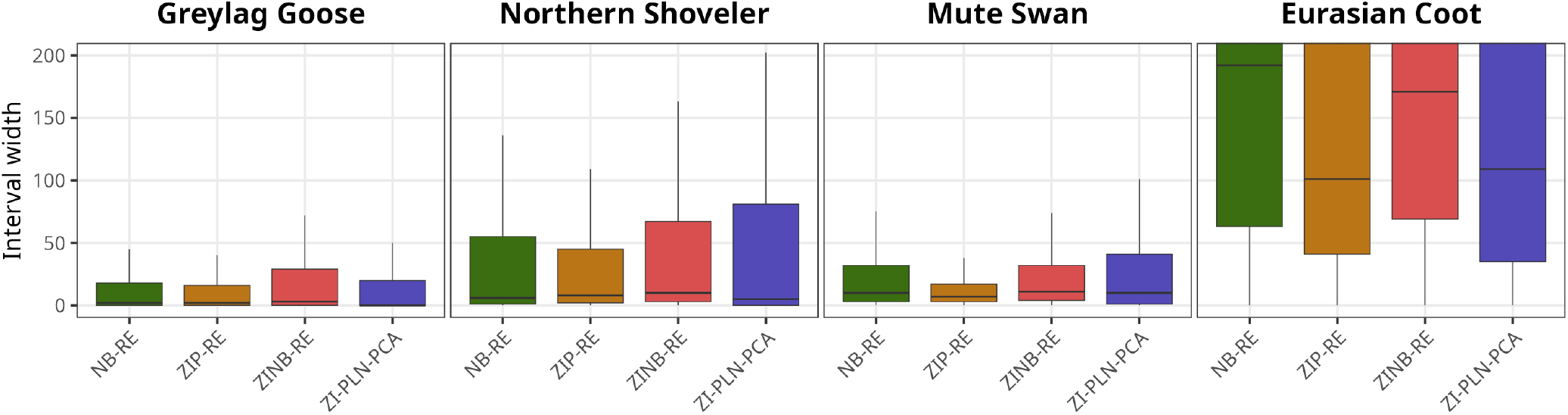
Width of prediction intervals for all species all rates of missing data combined for four models : NB-RE, ZIP-RE, ZINB-RE, ZI-PLN-PCA.

We saw in Table 4 that, in practice, we are unable to calculate 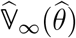 (using vcov for GLMMs). Thus, for certain models, there were too few replicates with a usable 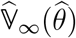 to calculate prediction intervals: the ZIP-FE, ZIP-MI, ZINB-FE and ZINB-MI models. We therefore only present the results for these models in Appendix A.3 Table 6.

**Table 6.**
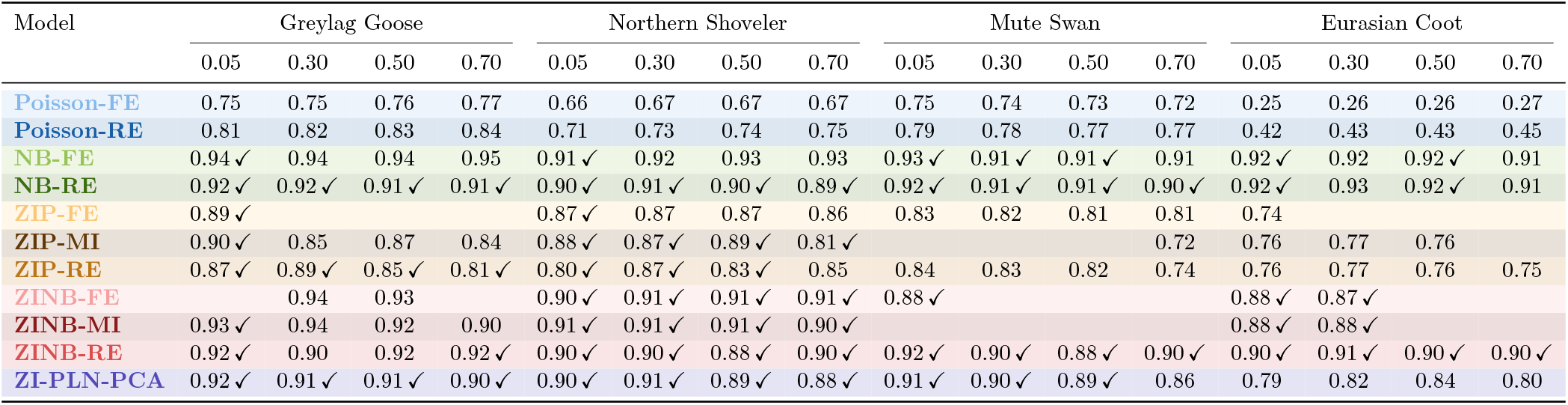
Mean empirical coverage rate, by species and missing-data rate. A ✓indicates that at least half of the replicates in that group have a coverage rate falling within their own binomial reference band Binomial(*M*, 0.9), where *M* is the number of missing observations for that replicate. Empty cells indicate configurations with no available replicates.

Among the remaining models, Poisson-FE and Poisson-RE almost never achieve the nominal coverage rate, with almost none of the replicates falling within the binomial bands across all species. NB-FE performs only slightly better for the Mute Swan alone. These models therefore appear unsuitable for species whose distribution combines zero-inflation and overdispersion.

Among the other models, all struggle to achieve the correct coverage rate for the Eurasian Coot. Either they over-cover (NB-FE and NB-RE) or they under-cover (ZIP-RE and ZI-PLN-PCA), except for ZINB-RE (see Table 5). This is because this species has the highest variance, and ZINB-RE is a model with two sources of overdispersion – the latent Gamma variable and random site effects – and it also models zero-inflation, which NB-RE does not. It is therefore the most likely to be able to account for both zeros and very high values. For rarer species like the Greylag Goose, the ZIP-RE and ZI-PLN-PCA models are the most suitable, ZINB-RE tend to over-cover in that case. But, ZIP-RE is not at all suitable for the Mute Swan, a moderately common species in the region, unlike ZI-PLN-PCA.

As regards the widths of the prediction intervals, we present only those for the four models that most consistently achieve the nominal coverage rate across species. For these four models, the widths of the intervals are consistent with the orders of magnitude of the counts for the various species (see Figure 1). The intervals are at most of the order of a hundred for the Greylag Goose, Mute Swan and Northern Shoveler, but of the order of a thousand for the Eurasian Coot (see Appendix A.3, Figure 15).

Overall, for rare to moderately common species (Greylag Goose, Northern Shoveler, Mute Swan), the ZI-PLN-PCA model most often achieves the nominal coverage rate with interval widths equivalent to those of the other models. However, for widespread and generalist species forming very large groups such as the Eurasian Coot, its intervals are too narrow to achieve the nominal coverage rate and ZINB-RE is more suitable.

##### Predictions of zeros

We now turn to the ability of each method to predict null counts. As explained in Section 3.1.4, we see this problem as a classification problem, that is being right at predicting a zero or not. Figure 6 compares the sensitivity (as defined in (16)) of each method for the four species. Because the results are similar for any missing data rate, all rates are pooled in the figure. The results for the balanced accuracy and for the AUC are given in Appendix A.4, Figures 16 and 17, respectively.

**Figure 6.**
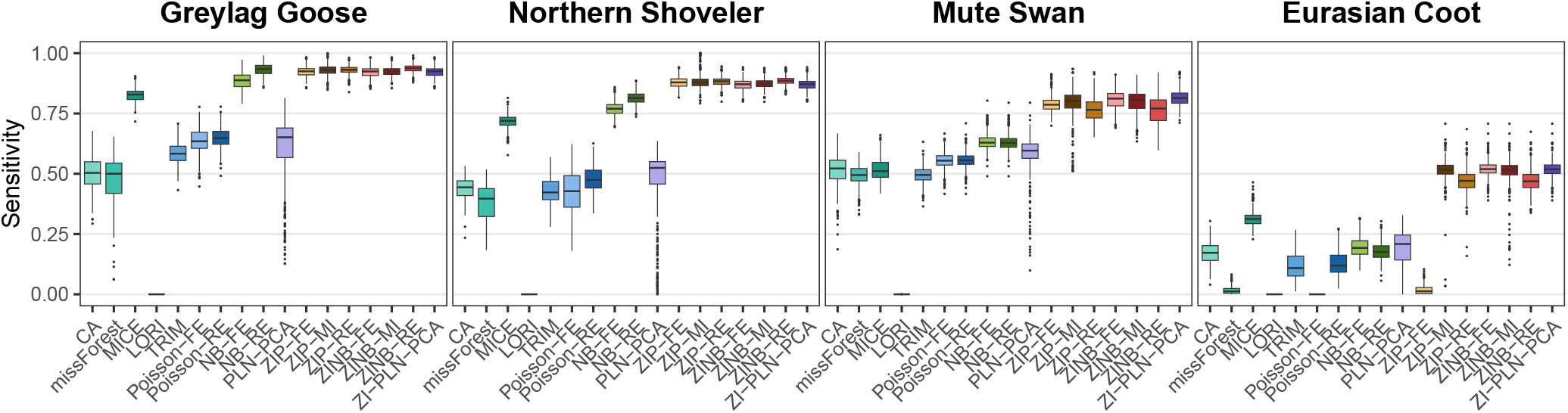
Sensitivity of zero classification for each species (all missing rates pooled).

The four species have different distribution (see Figure 1) and different proportion of null counts in their distribution. The Greylag Goose has the highest proportion of zeros and the Eurasian Coot the lowest. For all methods, we can see that the smaller the proportion of zero is, the harder it is to predict them.

A first observation is that GLMs, GLMMs, and ZI-PLN-PCA achieve markedly higher sensitivity, balanced accuracy, and AUC than the remaining methods. Among these statistical models, those that explicitly model presence (ZIP, ZINB, ZI-PLN-PCA) are best at predicting zeros, as might be expected, particularly when the proportion of zeros decreases – for the Eurasian Coot, for example: the Poisson model fails to capture the excess of zeros beyond what its mean-variance relationship predicts, and whilst the Negative Binomial model partially compensates through overdispersion, only zero-inflated models are truly suited to capturing this additional source of zeros, and thus predicting the 0s.

Three methods warrant specific attention. LORI’s sensitivity is consistently zero: not only does it fail to predict small values, as noted in the previous section, but it also fails to predict any zeros at all. Regarding Poisson-TRIM, this method automatically removes rows containing only zeros and missing data. We have therefore arbitrarily imputed zeros for these rows, which misrepresents its true ability to predict zeros. Finally, MICE should not be assessed on sensitivity alone: whilst its median sensitivity ranks in the middle of the field, its balanced accuracy is among the lowest. This is because MICE imputes based on neighboring observed values and, given the large proportion of zeros in these datasets, it naturally produces fewer errors on zero predictions whilst performing poorly on non-zero counts.

#### 3.2.3 Population estimate

So far, the comparison has focused on point-wise imputation errors. However, a common objective in ecological monitoring is to estimate population size, which typically relies on annual counts aggregated across sites. Moreover, these annual counts are used when one wants to estimate a temporal trend with a contrast method (see Section 2). Figure 7 therefore examines the difference between annual sums computed from imputed data and those computed from the complete dataset, for the Northern Shoveler across all missing data rates. Results for the other species (see Appendix A.5 in Figure 18) show similar results.

**Figure 7.**
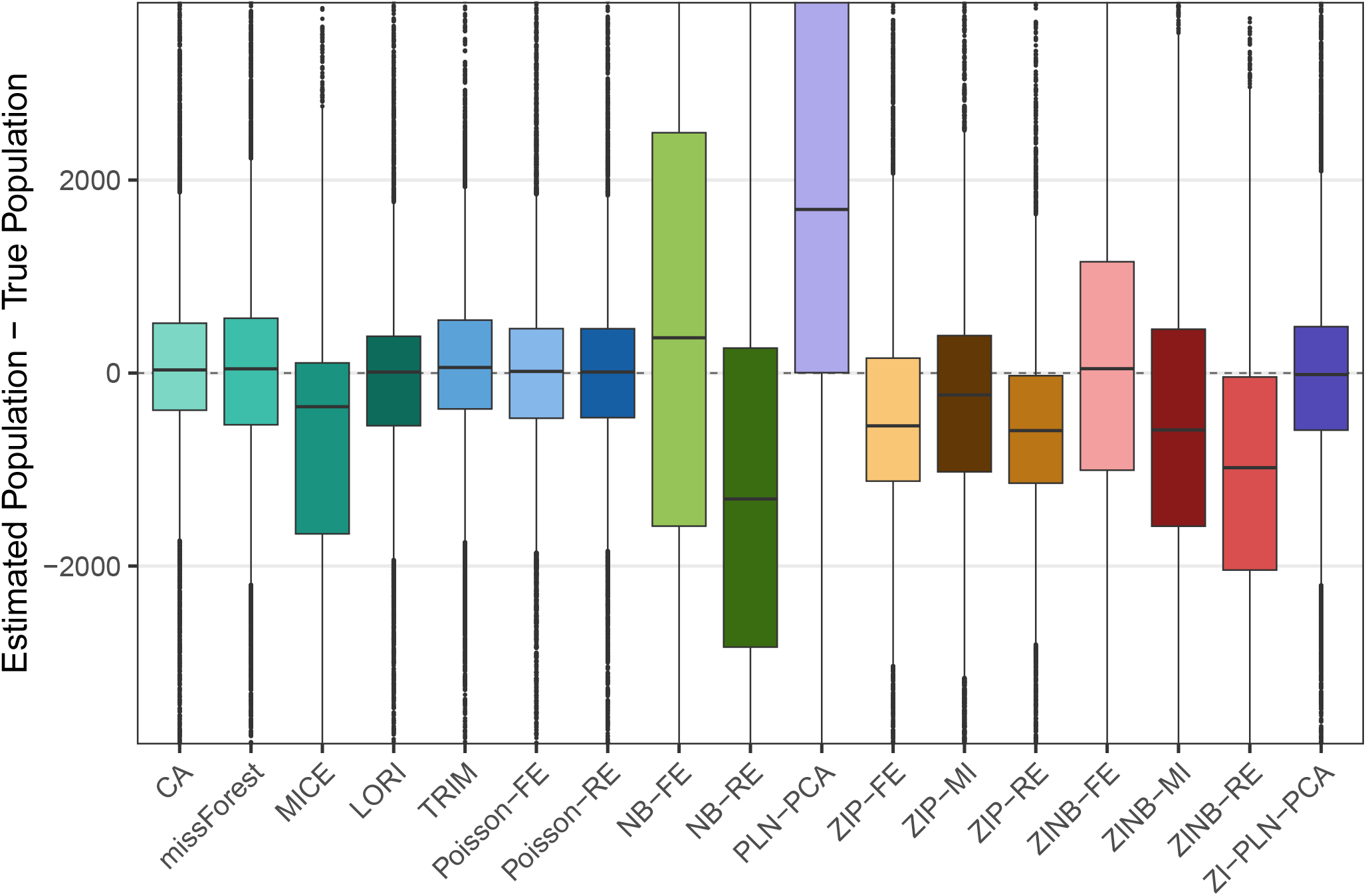
Differences between estimated populations (imputed annual sums) and true populations (true annual sums) for the Northern Shoveler, all missing rates. A point is equivalent to the sum of the abundances at all sites for a given year.

A first observation is that methods which perform modestly in terms of point-wise errors can prove entirely adequate for estimating annual sums. In particular, LORI, Poisson-TRIM, and the Poisson models all yield competitive results in this context. More generally, in absolute terms, the methods whose estimated populations are closest to the actual populations are CA, missForest, LORI, Poisson-TRIM, Poisson-FE, Poisson-RE and ZI-PLN-PCA.

Furthermore, these methods produce errors in population sizes centered around 0. The other methods systematically overestimate (PLN-PCA, NB-FE) or underestimate the population (MICE, NB-RE, ZIP’s and ZINB’s). This result is to be expected for GLMMs (NB-RE, ZIP-MI, ZIP-RE, ZINB-MI, ZINB-RE), as it was pointed out in Section 2.4 that the imputations used for these methods (the plug-in imputations) were consistently smaller than conditional imputations. This bias is evident in the estimate of the population size.

## 4 Discussion

Several conclusions can be drawn from this comparative evaluation. All the methods except from MICE and PLN-PCA achieve acceptable imputation quality, each with its own strengths depending on the criterion of interest. No single method dominates across all dimensions, but there are more advantages using model-based methods than contrast methods.

Comparing contrast methods (LORI, missForest, CA) with model-based methods (Poisson-TRIM, GLMs, GLMMs, PLN), we find that both provide equivalent point estimates, but the contrast methods do not provide a measure of uncertainty, such as prediction intervals around these estimates. And if we wish to go a step further and estimate a trend, we will not be able to account for the uncertainty caused by missing data around the trend when using contrast methods.

Apart from point estimates, certain methods prove to be more accurate for estimating population size, which is equivalent to the sum of all available sites for a given year: CA, missForest, LORI, Poisson-FE, Poisson-RE and ZI-PLN-PCA.

When we compare only the statistical models with one another, GLMs and GLMMs are much faster than PLN-PCA and ZI-PLN-PCA, but most suffer from convergence issues – particularly the zero-inflated models – which makes it impossible to compute 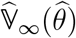. And in the context of these abundance data, even though point estimates are equivalent across all models, based on other criteria (prediction of zero and coverage with prediction intervals), it is clear that models which account for overdispersion and zero-inflation are more appropriate, particularly if one relies on the prediction of 0 or the coverage rate of the prediction intervals.

Finally, the appropriate model also depends on the specific abundance distribution of each species, and in particular its variance. While ZI-PLN-PCA and ZINB-RE are equivalent in terms of prediction intervals for the Mute Swan and the Northern Shoveler, the Eurasian Coot, which has by far the highest mean and variance among the four species, is better modelled by ZINB-RE, which contains two latent variables responsible for overdispersion (compared with one for the other models) and a latent variable to model presence. ZI-PLN-PCA, by contrast, is better suited to species with lower and less variable abundances, such as the Greylag Goose in France and Italy.

## Author Contribution

B.B. L.D. P.D.d.R. and T.G. conceived the ideas. B.B. S.D. and S.R. designed the methodology and led the writing of the manuscript. L.D. P.D.d.R. and T.G. ensured the acquisition of funding. All authors contributed critically to the drafts and gave final approval for publication.

## Data Availability

The data used in this study are subject to a data use agreement with the coordinator of the national programs responsible for collecting these data at the national level. The datasets, anonymised with respect to species names, analysed during the current study are available in the https://doi.org/10.6084/m9.figshare.33406693 repository.

## Acknowledgements

The authors would also like to thank Khalil Baddour (Tour du Valat) for extracting and formatting the datasets. We are grateful to the INRAE MIGALE bioinformatics facility (MIGALE, INRAE, 2020. Migale Bioinformatics Facility, doi: 10.15454/1.5572390655343293E12) for providing help and/or computing and/or storage resources. Many thanks to Mr. Marco Zenatello and Dr. Nicola Baccetti, Italian National Coordinators for ISPRA (Istituto Superiore per la Protezione e la Ricerca Ambientale), the ONCFS/FNC/FDC Wetlands Waterbird Network who collected the data.

## A Additional material

### A.1 Models

Here is a list of the models discussed in the article. It should be noted that *Y* is a matrix containing the abundances of a species across *n* sites and *p* years; *Y*_*ij*_ corresponds to the number of individuals of a species counted at site *i* in year *j. x* is a matrix containing *d* covariates, each with *n* × *p* coordinates (one coordinate per site-year pair); *x*_*ij*_ is a vector containing the coordinates of all the covariates for site *i* and year *j*.

**Poisson - Fixed Effects (Poisson-FE)**

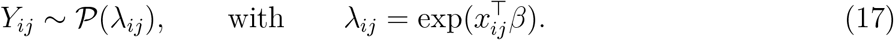

**Negative Binomial - Fixed Effects (NB-FE)**

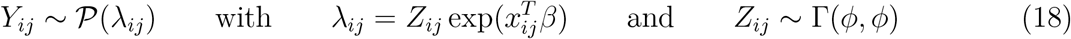

**Poisson Log-normal with dimensionality reduction (PLN-PCA)**

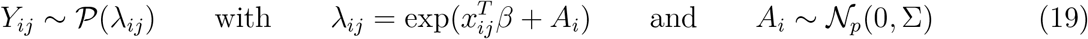

**Poisson - Random Effects (Poisson-RE)**

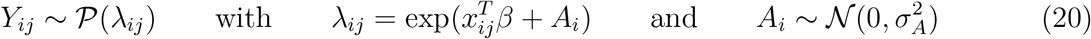

**Negative Binomial - Random Effects (NB-RE)**

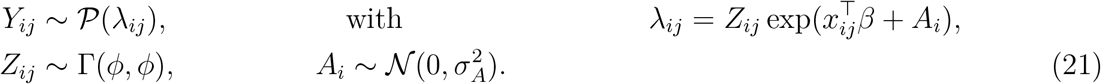

**Zero-inflated Poisson - Fixed Effects (ZIP-FE)**

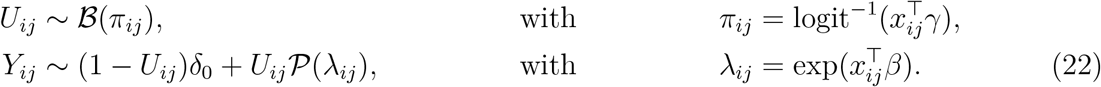

**Zero-inflated Negative Binomial - Fixed Effects (ZINB-FE)**

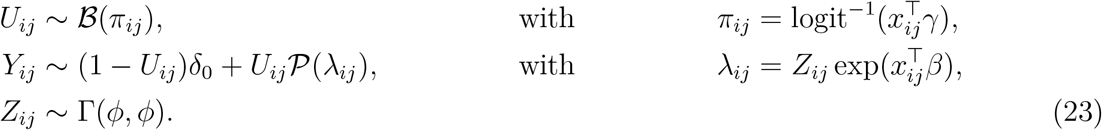

**Zero-inflated Poisson - Mixed Effects (ZIP-MI)**

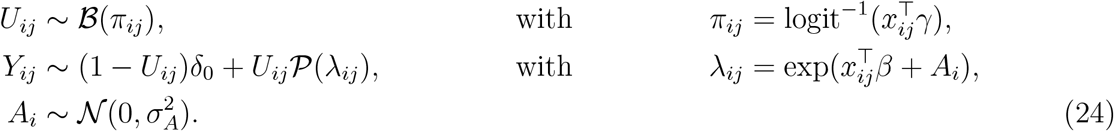

**Zero-inflated Negative Binomial - Mixed Effects (ZINB-MI)**

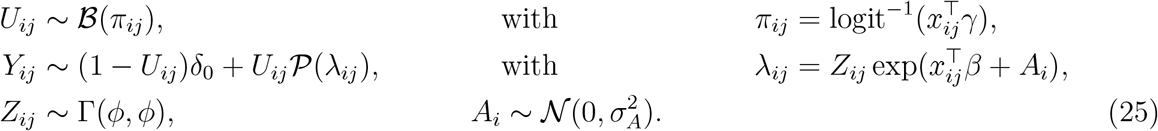

**Zero-inflated Poisson - Random Effects (ZIP-RE)**

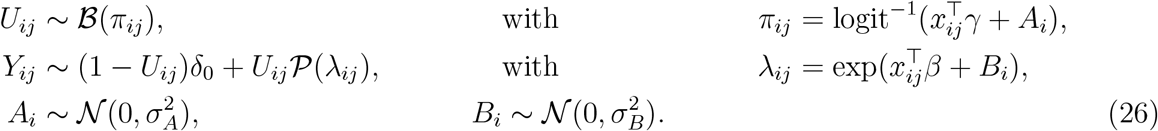

**Zero-inflated Negative Binomial - Random Effects (ZINB-RE)**

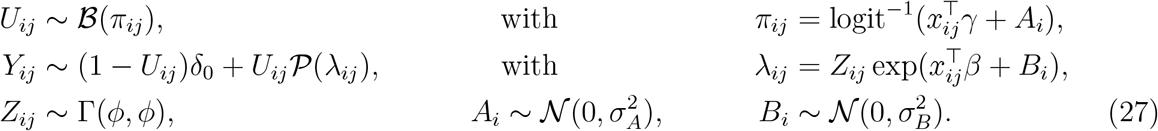

**Zero-inflated Poisson log-normal with dimensionality reduction (ZI-PLN-PCA)**

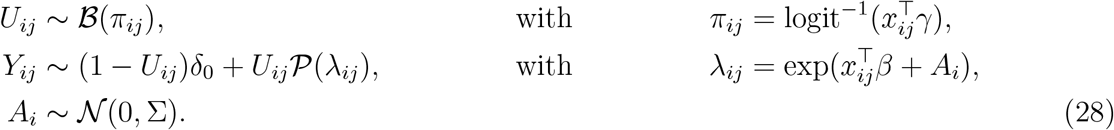

Σ has a low rank *q < p*, meaning there is a *p* × *q* matrix *B* such that Σ = *BB*^*T*^ .

### A.2 Imputations errors criteria

**Figure 8.**
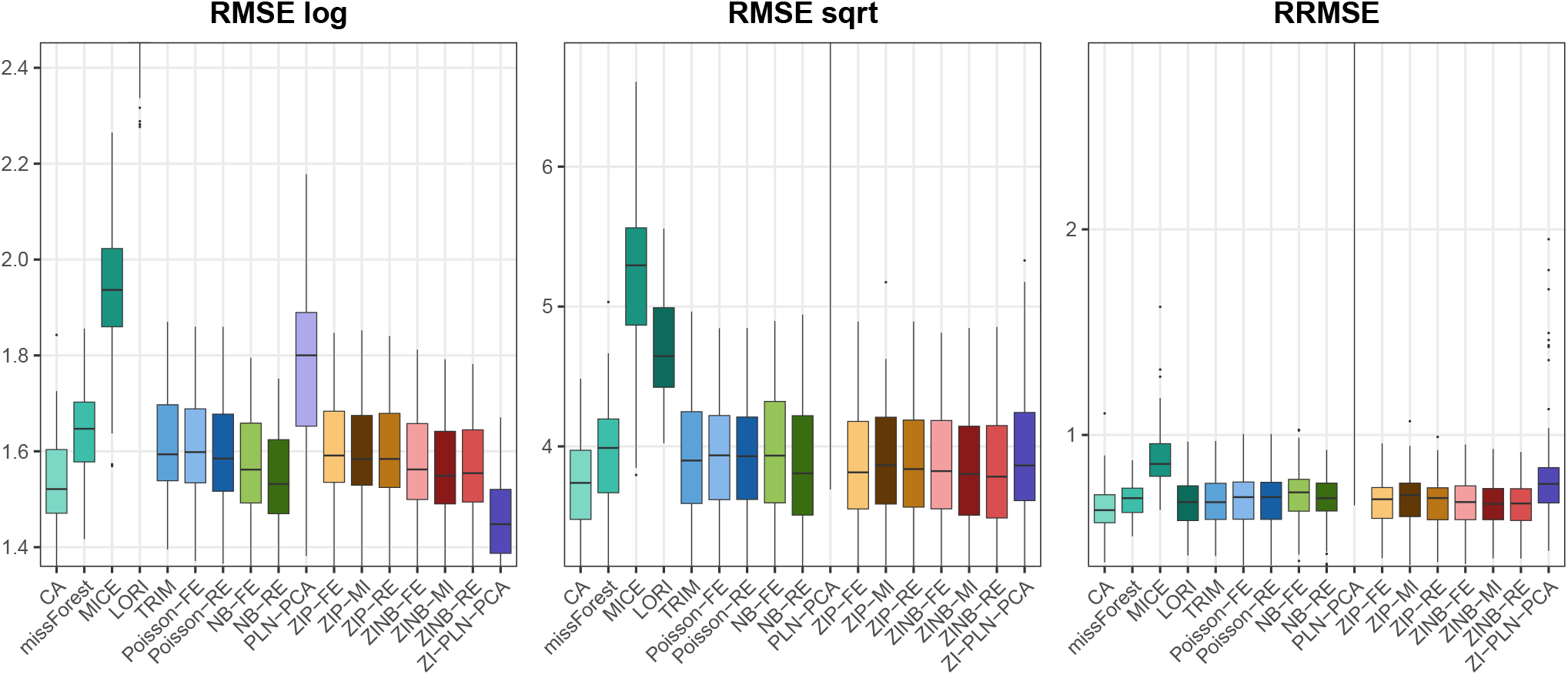
Imputation errors with 5% missing data for the northern shoveler for all methods measured using three different metrics.

**Figure 9.**
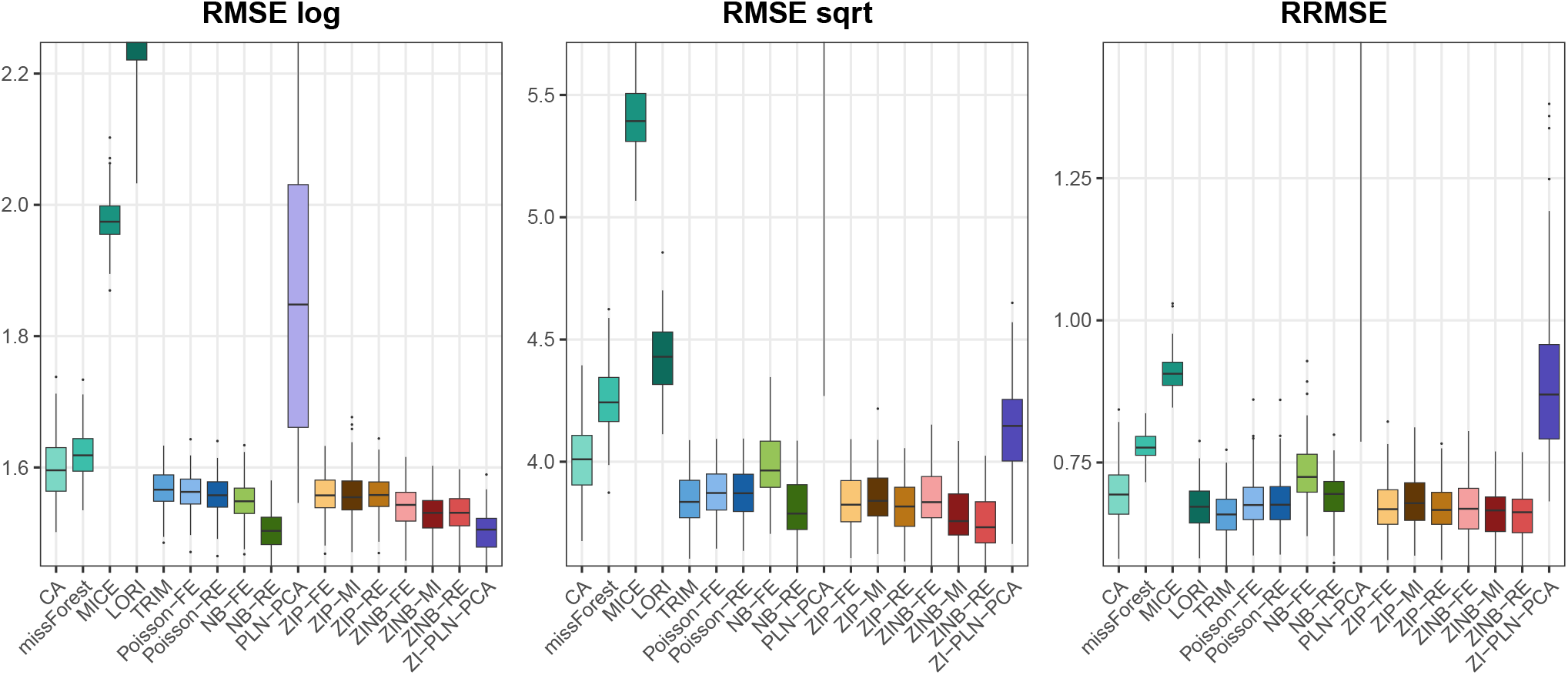
Imputation errors with 50% missing data for the northern shoveler for all methods measured using three different metrics.

**Figure 10.**
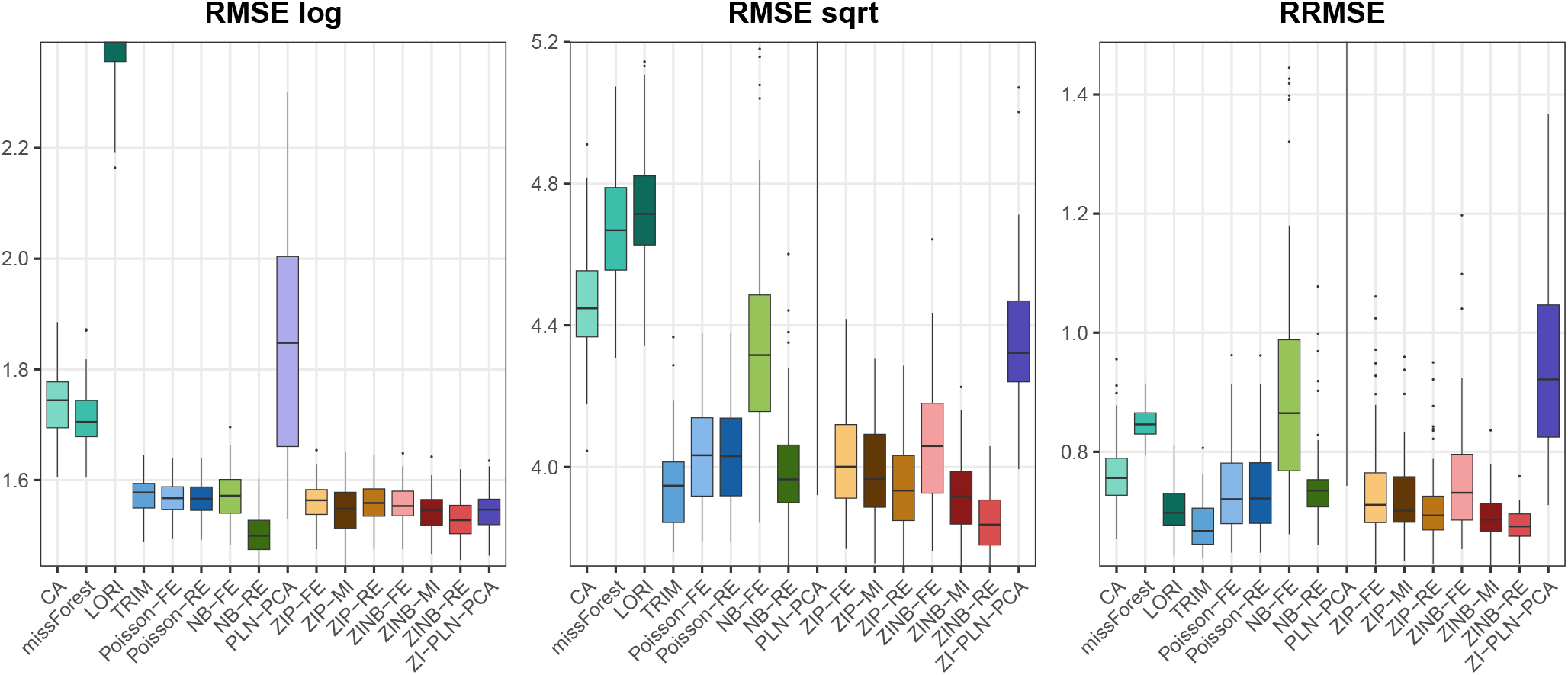
Imputation errors with 70% missing data for the northern shoveler for all methods measured using three different metrics.

**Figure 11.**
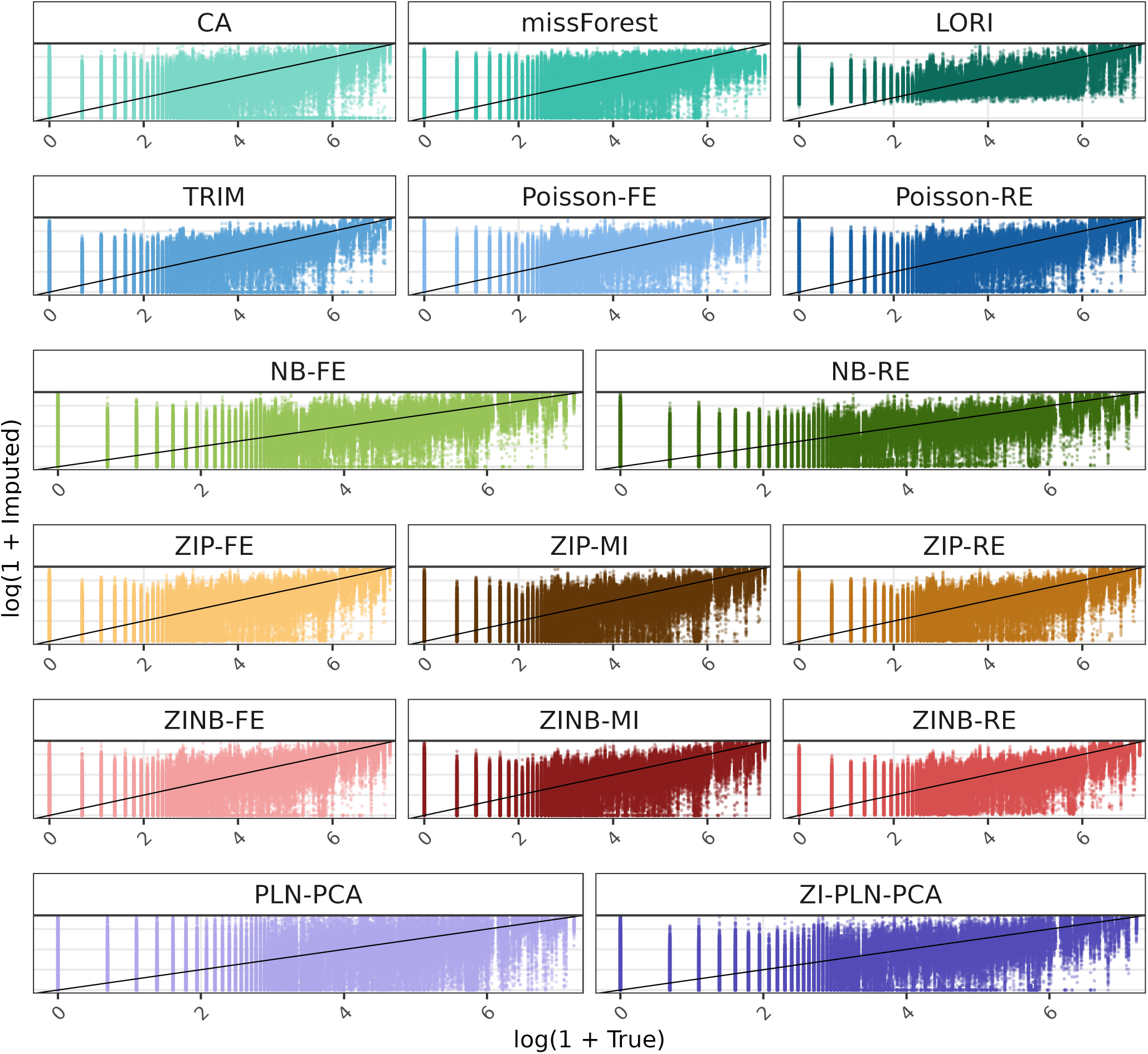
log(1 + Imputed values) with 70% missing data for the Northern Shoveler across all replicates versus log(1 + truth) for each method.

### A.3 Prediction intervals

#### A.3.1 Estimation of prediction intervals

Here we present the two algorithms used to estimate prediction intervals using the Monte Carlo method. There are two different algorithms because there are several types of prediction (see Section 2.4): marginal predictions, conditional predictions and so-called plug-in predictions. Algorithm 1 describes the estimated prediction intervals for plug-in predictions, used for the GLMMs, and algorithm 2 describes the estimated prediction intervals for conditional predictions, used for ZI-PLN-PCA.

##### Algorithm 1

Plug-in confidence and prediction intervals (Monte Carlo) for random-effects GLMMs (here Poisson-RE)

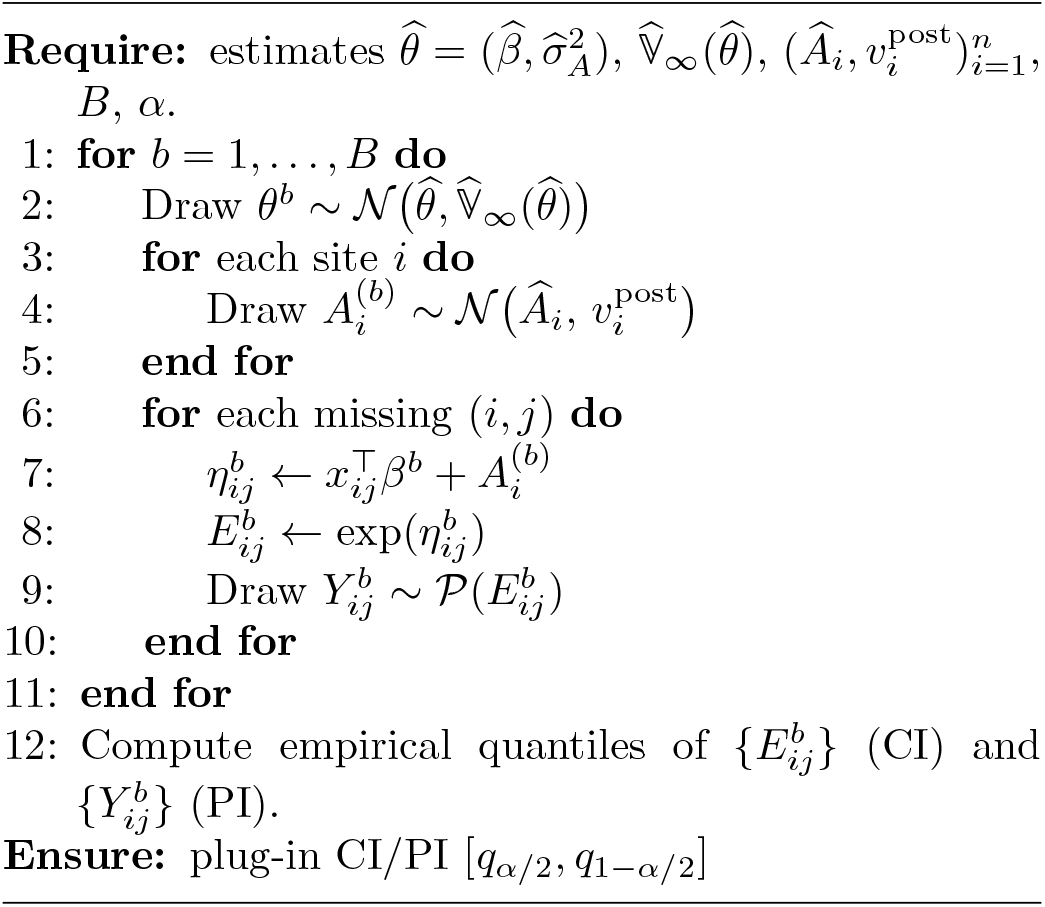

##### Algorithm 2

Conditional confidence and prediction intervals (Monte Carlo) for ZI-PLN-PCA

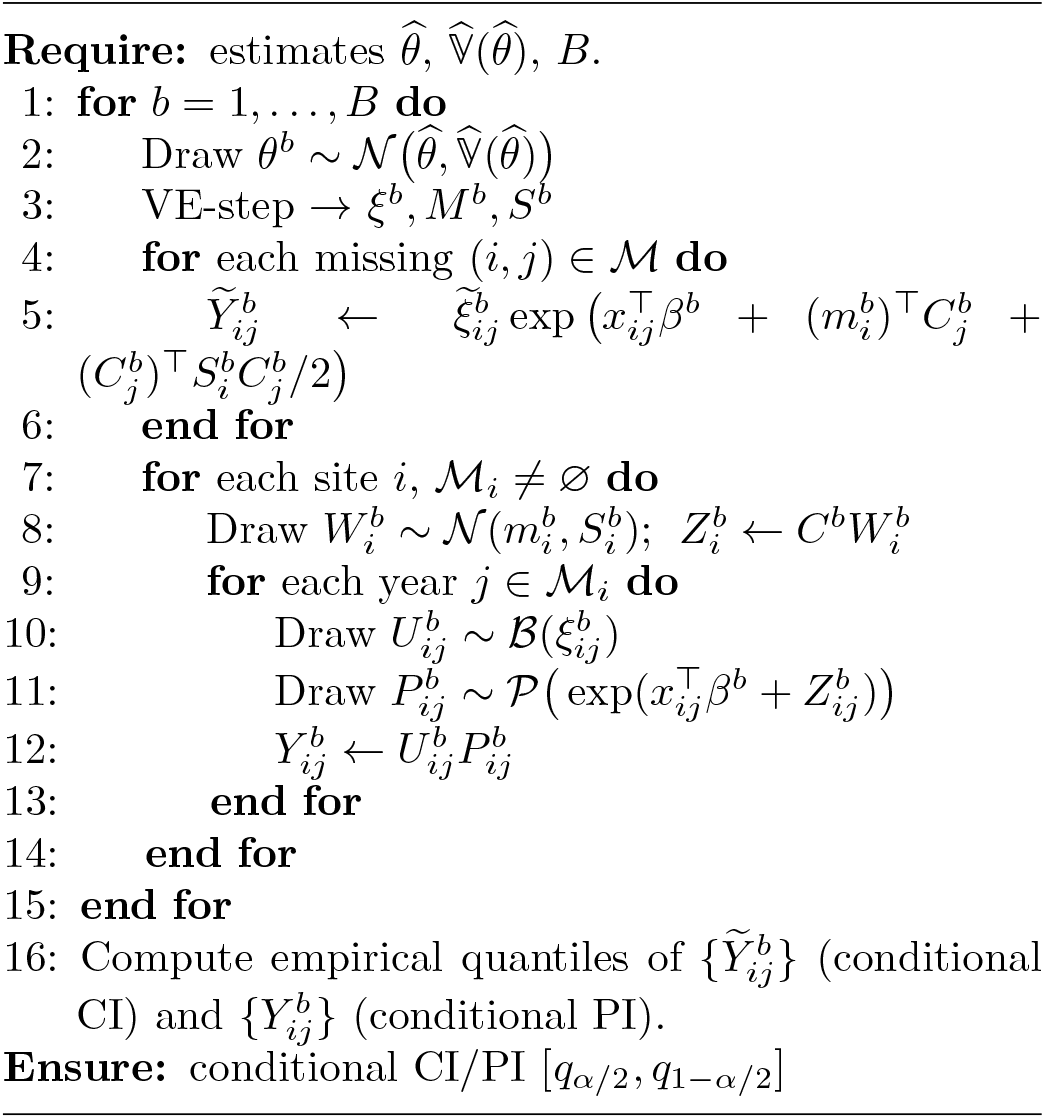

Note that the difference between the two algorithms lies in the third step. In algorithm 2, a VE step is performed to resample the variational parameters, whereas this is not done in algorithm 1. Instead, the BLUPs are drawn from a distribution estimated using the Laplace approximation. Furthermore, for all models without random effects (GLMs), where marginal predictions are used (as conditional predictions are meaningless in this case), either of these two algorithms can be used without having to carry out the third step.

Finally, the algorithm 2 has been developed to calculate the prediction intervals for the ZI-PLN-PCA model, and algorithm 1 is a basic algorithm for GLMMs, which corresponds to Poisson-RE, but it can be modified to suit other GLMMs. In particular, as in algorithm 2, steps 10 and 12 can be reused for zero-inflated models (ZIP-FE, ZIP-MI, ZINB-FE, ZINB-MI). If there are random effects in the presence part of the model (ZIP-RE, ZINB-RE), one must proceed in the same way as for the abundance part and obtain the distributions of the BLUPs estimated by the Laplace approximation. Finally, the algorithm 1 can also be adapted to negative binomial models.

#### A.3.2 Coverage rate of the prediction interval

We present the coverage rates for all species, and rates of missing data.

When a cell is empty, this means that we were unable to find 10 replicates out of 100 for which we could calculate 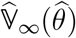 using vcov.

#### A.3.3 Prediction interval width

We present the prediction interval widths for all the species and all the models that have prediction intervals.

**Figure 12.**
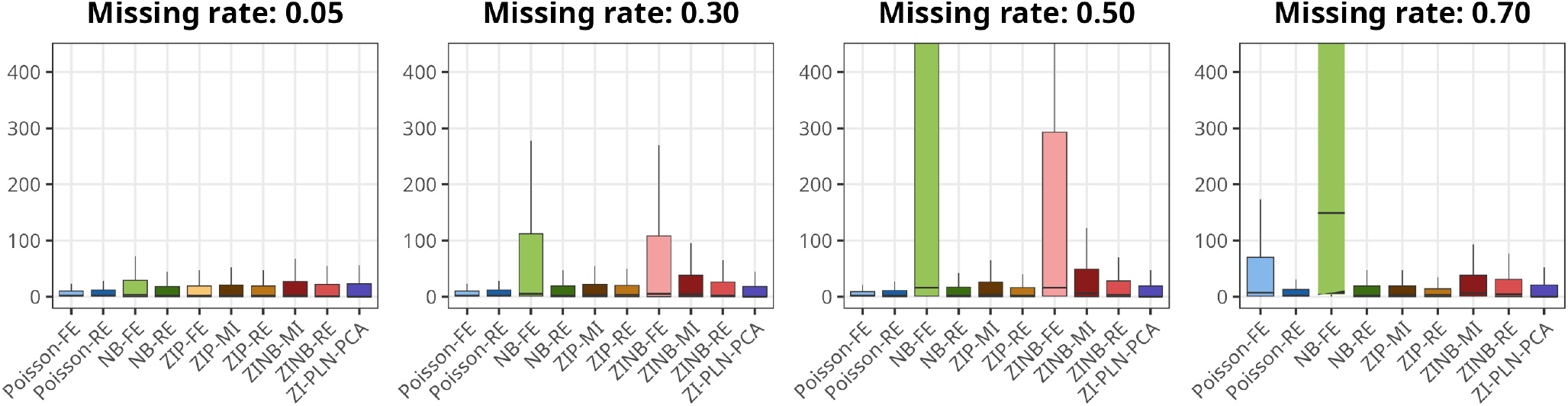
Prediction interval widths for the Greylag Goose.

**Figure 13.**
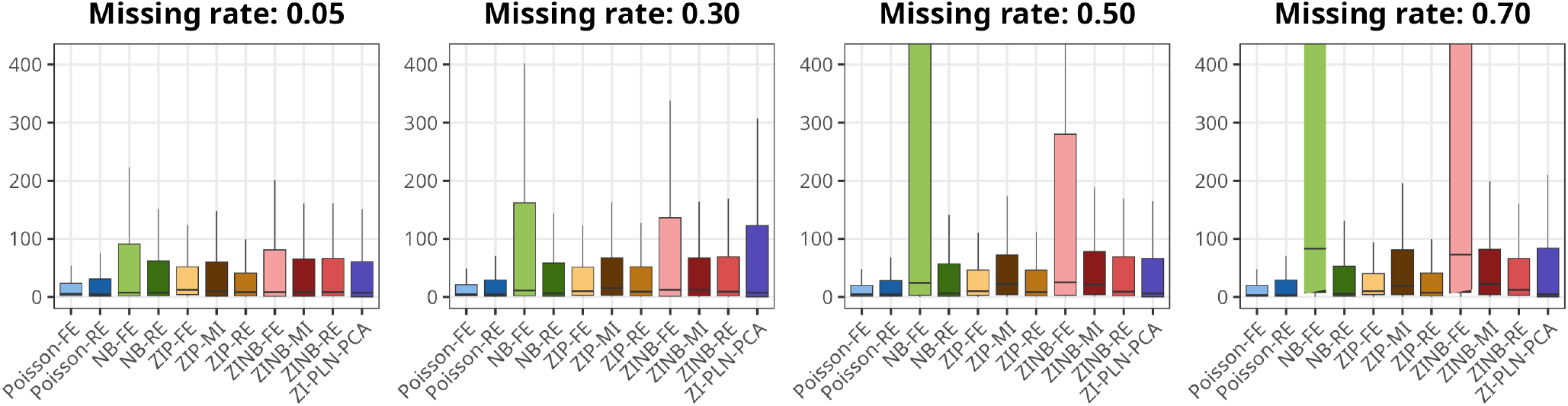
Prediction interval widths for the Northern Shoveler.

**Figure 14.**
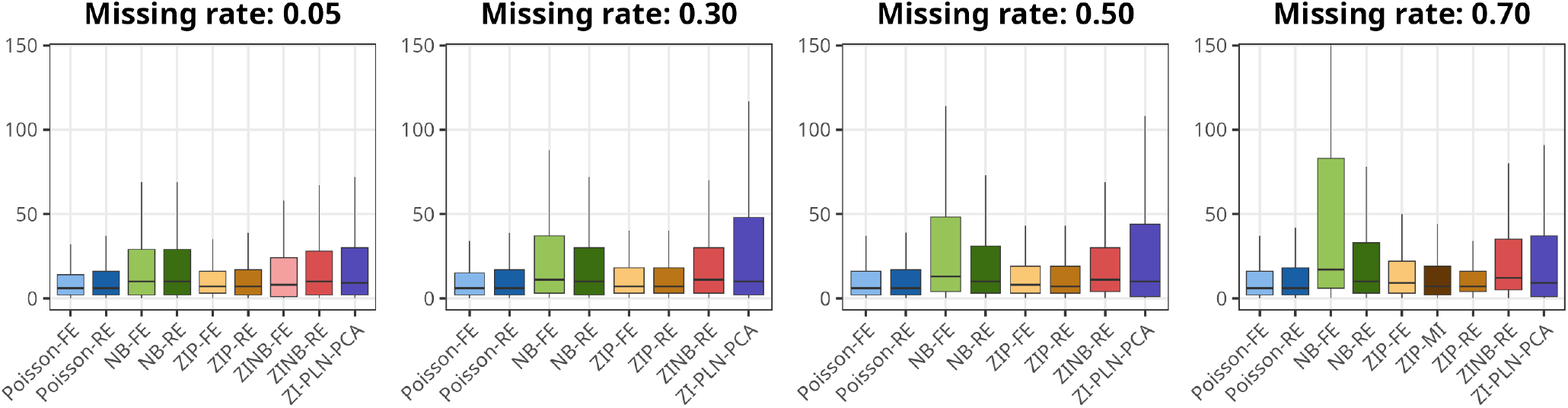
Prediction interval widths for the Mute Swan.

**Figure 15.**
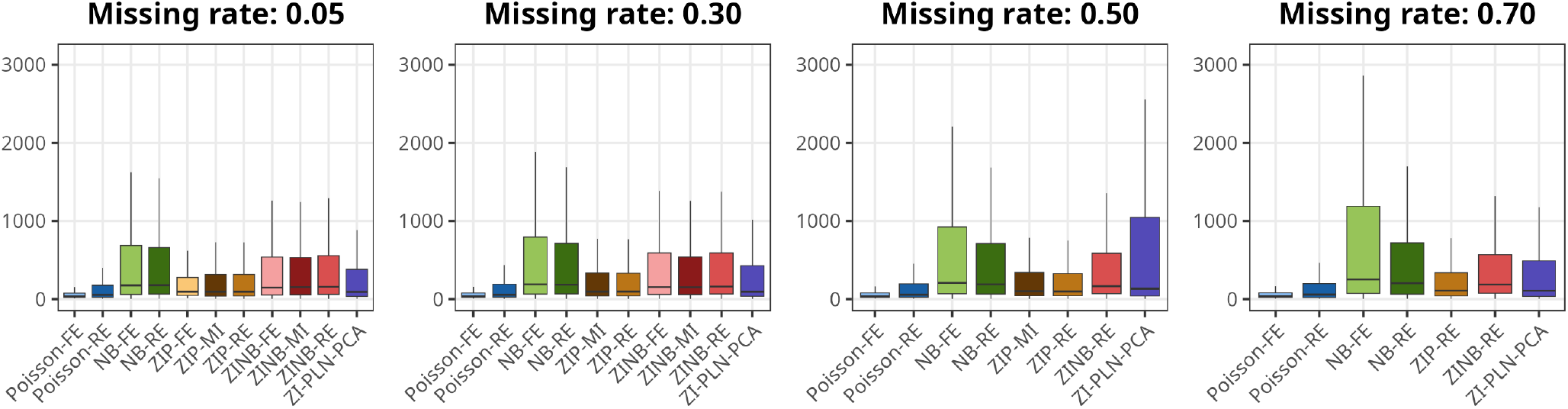
Prediction interval widths for the Eurasian Coot.

### A.4 Prediction of zeros

**Figure 16.**
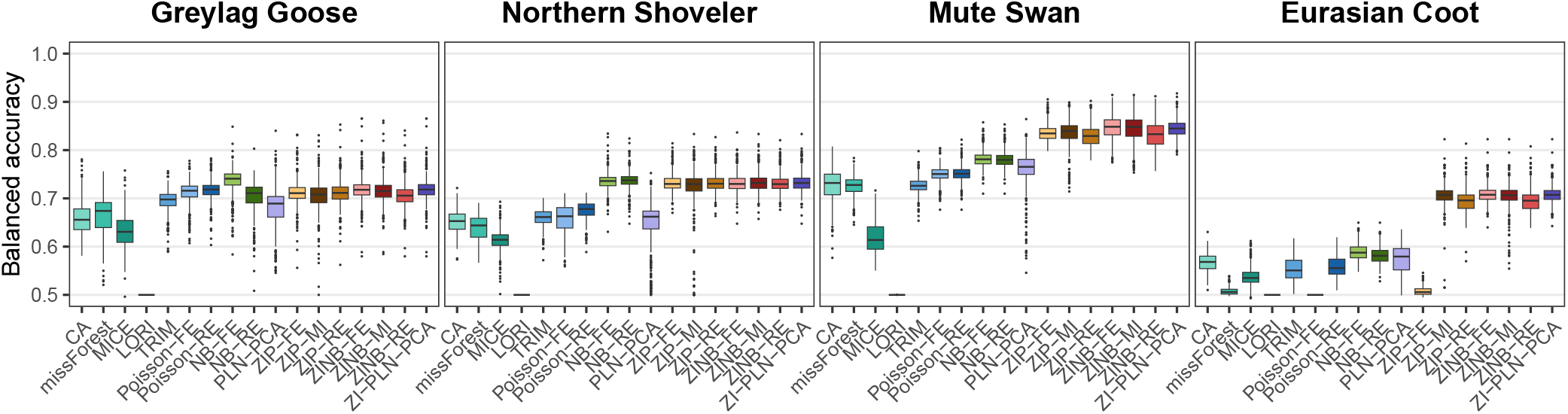
Balanced accuracy of zero classification for each species (all missing rates pooled).

**Figure 17.**
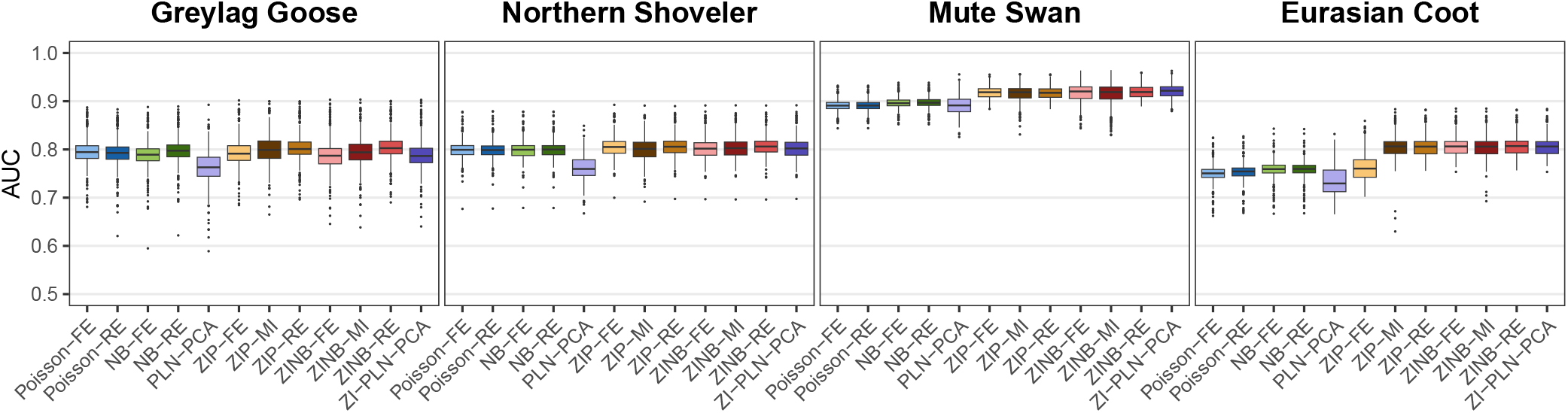
AUC of zero classification for each species (all missing rates pooled).

### A.5 Population estimation

**Figure 18.**
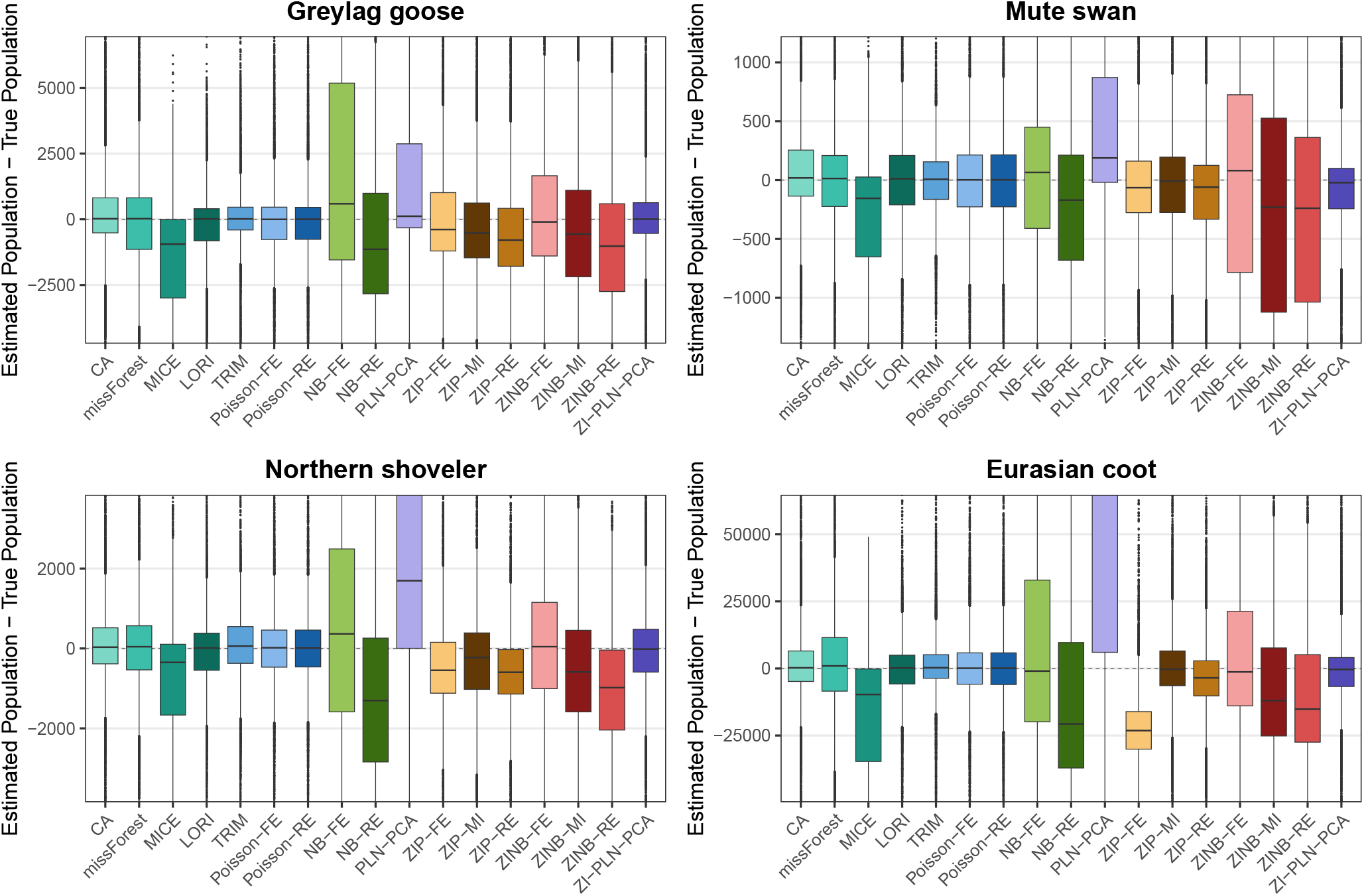
Differences between estimated population (imputed annual sums) and true population (true annual sums) for all species, all missing rates pooled.

